# Functional Communication between IP_3_R and STIM2 at sub-threshold stimuli is a critical checkpoint for initiation of SOCE

**DOI:** 10.1101/2021.09.24.461545

**Authors:** Moaz Ahmad, Hwei Ling Ong, Hassan Saadi, Ga Yeon Son, Zahra Shokatian, Lara E. Terry, Mohamed Trebak, David I. Yule, Indu Ambudkar

## Abstract

Stromal interaction molecules, STIM1 and STIM2, sense decreases in the endoplasmic reticulum (ER) [Ca^2+^] ([Ca^2+^]_ER_) and cluster in ER-plasma membrane (ER-PM) junctions where they recruit and activate Orai1. While STIM1 responds when [Ca^2+^]_ER_ is relatively low, STIM2 displays constitutive clustering in the junctions and is suggested to regulate basal Ca^2+^ entry. The cellular cues that determine STIM2 clustering under basal conditions is not known. By using gene editing to fluorescently tag endogenous STIM2, we report that endogenous STIM2 is constitutively localized in mobile and immobile clusters. The latter associate with ER-PM junctions and recruit Orai1 under basal conditions. Agonist stimulation increases immobile STIM2 clusters which co- ordinate recruitment of Orai1 and STIM1 to the junctions. Extended synaptotagmin (E-Syt)2/3 are required for forming the ER-PM junctions, but are not sufficient for STIM2 clustering. Importantly, inositol 1,4,5-triphosphate receptor (IP_3_R) function and local [Ca^2+^]_ER_ are the main drivers of immobile STIM2 clusters. Enhancing, or decreasing, IP_3_R function at ambient [IP_3_] causes corresponding increase, or attenuation, of immobile STIM2 clusters. We show that immobile STIM2 clusters denote decreases in local [Ca^2+^]_ER_ mediated by IP_3_R that is sensed by the STIM2-N terminus. Finally, under basal conditions, ambient PIP_2_-PLC activity of the cell determines IP_3_R function, immobilization of STIM2, and basal Ca^2+^ entry while agonist stimulation augments these processes. Together, our findings reveal that immobilization of STIM2 clusters within ER-PM junctions, a first response to ER-Ca^2+^ store depletion, is facilitated by the juxtaposition of IP_3_R and marks a checkpoint for initiation of Ca^2+^ entry.

**Significance:** STIM proteins sense decreases in [Ca^2+^]_ER_ and cluster in endoplasmic reticulum (ER)-plasma membrane (PM) junctions where they recruit and activate Orai1. While STIM1 clustering requires substantial [Ca^2+^]_ER_ decrease, STIM2 displays pre-clustering under resting conditions and regulates basal Ca^2+^ entry. The mechanism(s) underlying constitutive clustering of STIM2 is not known. We show herein that endogenous STIM2 assembles as mobile and immobile clusters and that Orai1 is recruited to the latter. Anchoring of STIM2 clusters is triggered by decreases in local [Ca^2+^]_ER_ that is mediated by ambient activity of IP_3_R and sensed by the STIM2 N-terminus. This functional link between IP_3_R and STIM2 governs constitutive STIM2 clustering and ensures coupling of [Ca^2+^]_ER_ decrease at sub-threshold stimuli with activation of Ca^2+^ entry.

## Introduction

Store-operated calcium entry (SOCE), which provides critical cytosolic Ca^2+^ signals for regulation of diverse cell functions, is activated in response to depletion of Ca^2+^ stores within the endoplasmic reticulum (ER) (1, 2). Decreases in [Ca^2+^]_ER_ are sensed by resident ER proteins <u>St</u>romal <u>I</u>nteraction <u>M</u>olecules 1 and 2 (STIM1 and STIM2) via their N-terminal Ca^2+^-binding domains localized in the ER lumen. This triggers oligomerization and clustering of STIM proteins in ER-plasma membrane (PM) junctions, (3–5) where they recruit and activate the PM channel Orai1, which mediates SOCE (6–10). STIM1, the primary regulator of Orai1, has a relatively high Ca^2+^ affinity and responds only when [Ca^2+^]_ER_ is substantially decreased. In contrast, STIM2, a relatively weak activator of Orai1, has low Ca^2+^ affinity which enables the protein to respond to minimal decreases in [Ca^2+^]_ER_ (4, 9–12). When overexpressed in cells, STIM2 displays constitutive clustering within ER-PM junctions. It also recruits and activates Orai1 channels in these junctions causing Ca^2+^ entry in unstimulated cells (4, 12, 13). Our previous findings demonstrate that pre-clustering of STIM2 promotes recruitment of Orai1/STIM1 and facilitates STIM1 activation under conditions when [Ca^2+^]_ER_ is not sufficiently depleted to activate STIM1 (9, 10). These previous data suggest that pre-clustered STIM2 is in an activated state in unstimulated cells and contributes to basal Ca^2+^ entry. There is, however, little information regarding the clustering of endogenous STIM2 and the molecular mechanisms, or cellular cues, that regulate its pre-clustering at ER-PM junctions in the cell. A particular concern is that exogenous overexpression of STIM2 alters the stoichiometry of endogenous STIM/Orai complexes, which might artificially force them into the junctions to cause pre-clustering.

In this study, we have examined endogenous STIM2 and its regulation under physiological conditions. We have used CRISPR/Cas9 to knock-in mVenus into the N-terminus of native *Stim2* gene, and generated HEK293 cell lines expressing fluorescently tagged endogenous STIM2 (mV- STIM2). We report that endogenous STIM2 is pre-clustered in the ER-PM junctional region of cells under basal conditions. While majority of STIM2 clusters are mobile, there is a small population of relatively immobile STIM2 clusters. Importantly, immobilization of native STIM2 clusters is triggered by decreases in local [Ca^2+^]_ER_ that are mediated by functional IP_3_ receptors (IP_3_R) and sensed by STIM2 N-terminus. In the absence of added agonist, constitutive PIP_2_-PLC activity, together with cAMP/protein kinase A (PKA) signaling, determines IP_3_R function. Consistent with this response of STIM2 at ambient stimuli, there is an increase in immobile STIM2 clusters following simulation of cells with a Ca^2+^-mobilizing agonist. Further, the immobile STIM2 clusters demarcate sites where Orai1 clusters in basal conditions and both Orai1 and STIM1 cluster following agonist stimulation. Together, our findings suggest that a critical functional link between IP_3_R and STIM2 underlies pre-clustering of STIM2 and is a checkpoint for initiation of SOCE in response to decreases in [Ca^2+^]_ER_.

## Results

### Endogenous STIM2 is pre-clustered in ER-PM junctions

mVenus was knocked in at the N-terminus of *Stim2* gene using the CRISPR/Cas9 approach in HEK293 cells (experimental details are provided in Fig. S1 *A* and *B*) and integration of mVenus was confirmed by PCR and genomic sequencing. HEK293 cells expressing mVenus-tagged endogenous STIM2 (mV-STIM2, indicated as S2 in all the figures) were isolated by single-cell sorting (Fig. S1 *C* and *D*) and used to establish a monoclonal cell population (STIM2-KI cells, which were used for all the experiments described below unless indicated otherwise). All STIM2-KI cells expressed mV-STIM2, which was substantially reduced by siSTIM2 (Fig. S1*E*). STIM2 expression was similar in wild-type HEK293 (WT) and STIM2-KI cells, note presence of mV-STIM2 in latter (Fig. S1*F*). Further, expressions of STIM1 and Orai1, were similar in WT and STIM2-KI cells. Cyclopiazonic acid (CPA)-induced internal Ca^2+^ release and Ca^2+^ entry were similar in WT and STIM2-KI cells (Fig. S1 *G* to *J*). In both sets of cells, knockdown of either STIM protein expression significantly reduced Ca^2+^ influx, with relatively greater decrease induced by STIM1 knockdown.

mV-STIM2 displayed a reticular pattern of localization in STIM2-KI cells (Fig. 1*A* confocal image; an enlarged view shown in the right panel). mV-STIM2 co-localized with an ER marker (mCherry-ER3, mCh-ER3) that was expressed in STIM2-KI cells (Fig. 1*B*, also see magnified images), However, mV-STIM2 signal was not uniform in the ER but appeared to be punctate (Pearson’s Correlation Coefficient (PCC) = 0.76 and Mander’s Overlay Coefficient (MOC) = 0.98/0.98). mV-STIM2 was detected as clusters in the TIRF plane in unstimulated cells (Fig. 1*C*). mCh-ER3 expressed in WT HEK293 cells was also detected in the this cellular region (Fig. 1*D*). mV-STIM2 was localized within this ER network in STIM2-KI cells expressing mCh-ER3 (Fig. 1*E,* enlarged area shown in Fig. 1*F*, also see line scans; PCC=0.57 and MOC =0.99/0.97). Although not all areas of ER had STIM2, every STIM2 cluster was co-localized with ER. A substantial number of endogenous mV-STIM2 clusters were recruited to ER-PM junctions, in the absence of ER-Ca^2+^ store depletion, when MAPPER (14) was expressed in STIM2-KI cells (mV- STIM2 was co-localized with mCerulean-MAPPER, PCC=0.75 and MOC M1/M2 = 0.89/0.97, line scans show overlapping peaks of both proteins (Fig. 1*G*)).

**Fig. 1.**
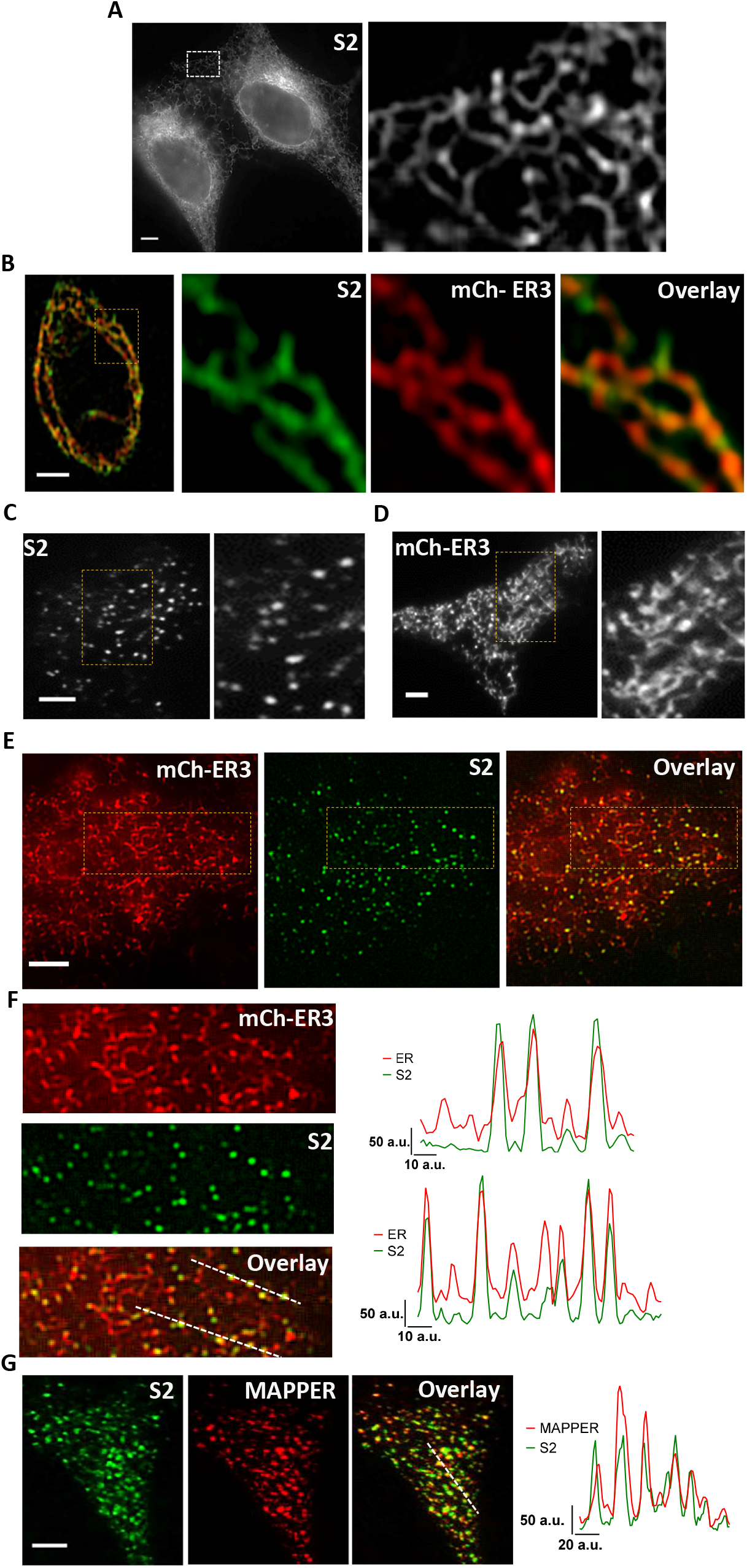
Visualization of endogenous STIM2. (*A*) Localization of endogenous mVenus-STIM2 (mV-STIM2; S2 in images, acquired by confocal microscopy) in STIM2-KI cells, enlargement of the demarcated area shown on right. (*B*) Airyscan images with enlarged region of STIM2-KI cell showing mV-STIM2 (green), expressed mCherry-ER3 (mCh-ER3; Red) and overlay. (*C*) TIRF images of STIM2-KI cells in basal, unstimulated conditions; enlargement of demarcated area on right. (*D*) Localization of mCh-ER3 in HEK293, marked area enlarged in right image. (*E*) Images from left to right: mCh-ER3 expressed in STIM2-KI cells (ER; Red), mV-STIM2 (S2; Green) and overlay. (*F*) Enlarged images of demarcated regions in *E* and line scans (position indicated in overlay image- mCh-ER3 (ER; Red) and mVenus-STIM2 (S2; Green)). (*G*) mCerulean-MAPPER (MAPPER, Red) expressed in STIM2-KI cells and mV-STIM2 (S2; Green), and overlay; line scans (position indicated in overlay image) show mCerulean-MAPPER (Red) and mV-STIM2 (Green). All microscope images show representative cells from at least 3 experiments. Scale bars, 5µm.

### Endogenous STIM2 exists as immobile and mobile clusters

When STIM2-KI cells were stimulated with relatively low (1μM) or high (100μM) carbachol (CCh), there was a time- and dose-dependent increase in the number and fluorescence intensity of mV-STIM2 clusters (Fig. 2 *A* and *B*, addition of the agonist at 2 min is indicated; Supplementary Videos 1 and 2). Activation-induced clustering of STIM1 and Orai1 in ER-PM junctions is associated with a decrease in their mobility (15). Since STIM2 is suggested to be in an activated state under basal conditions, we examined the mobility of mV-STIM2 in unstimulated cells by identifying those that remained at the same location for at least 1 min (similar criteria have been previously used to determine stability of proteins, including IP_3_R; (16)). A small proportion of mV-STIM2 clusters was relatively immobile under ambient conditions (Fig. 2 *C* and *D* show enlarged areas of STIM2-KI cells, whole cell image is shown in Fig. S2). The images in Fig. 2 *C* and *D* labeled as “basal” show mV-STIM2 clusters (green) present in the first frame (indicated as 0) overlaid with the image acquired 1 min later (0/1m, mV-STIM2 clusters in second frame are indicated magenta). Clusters that overlap (white) represent those that remained at the same location for 1 min (also shown by, line scans to the right of the image). Note that relatively immobile clusters displayed similar location and fluorescence amplitude in subsequent frames, while mobile clusters appeared as non-overlapping magenta and green peaks with different amplitudes. Importantly, the number of these immobile clusters was increased dose-dependently following stimulation of the cells with low (1µM) or high (100µM) CCh (Fig. 2 *C* and *D*; appear as white clusters in the images and as overlapping peaks in the line scans). Cells were stimulated at the 2 min time point, and overlay images from 3 and 4 min (3/4m; i.e. 1 and 2 min after stimulation) were used to identify early changes in mobility of mV-STIM2 clusters. Time-dependent changes in immobile mV-STIM2 clusters was quantified (Fig. 2*E,* using the Image Calculator tool in Fiji/Image2). The time- increase in immobile mV-STIM2 clusters (relative to the number of immobile clusters prior to stimulation) was higher at 100µM than 1µM CCh (also reflected by the increase in PCC values (Fig. 2*F*)).

**Fig. 2.**
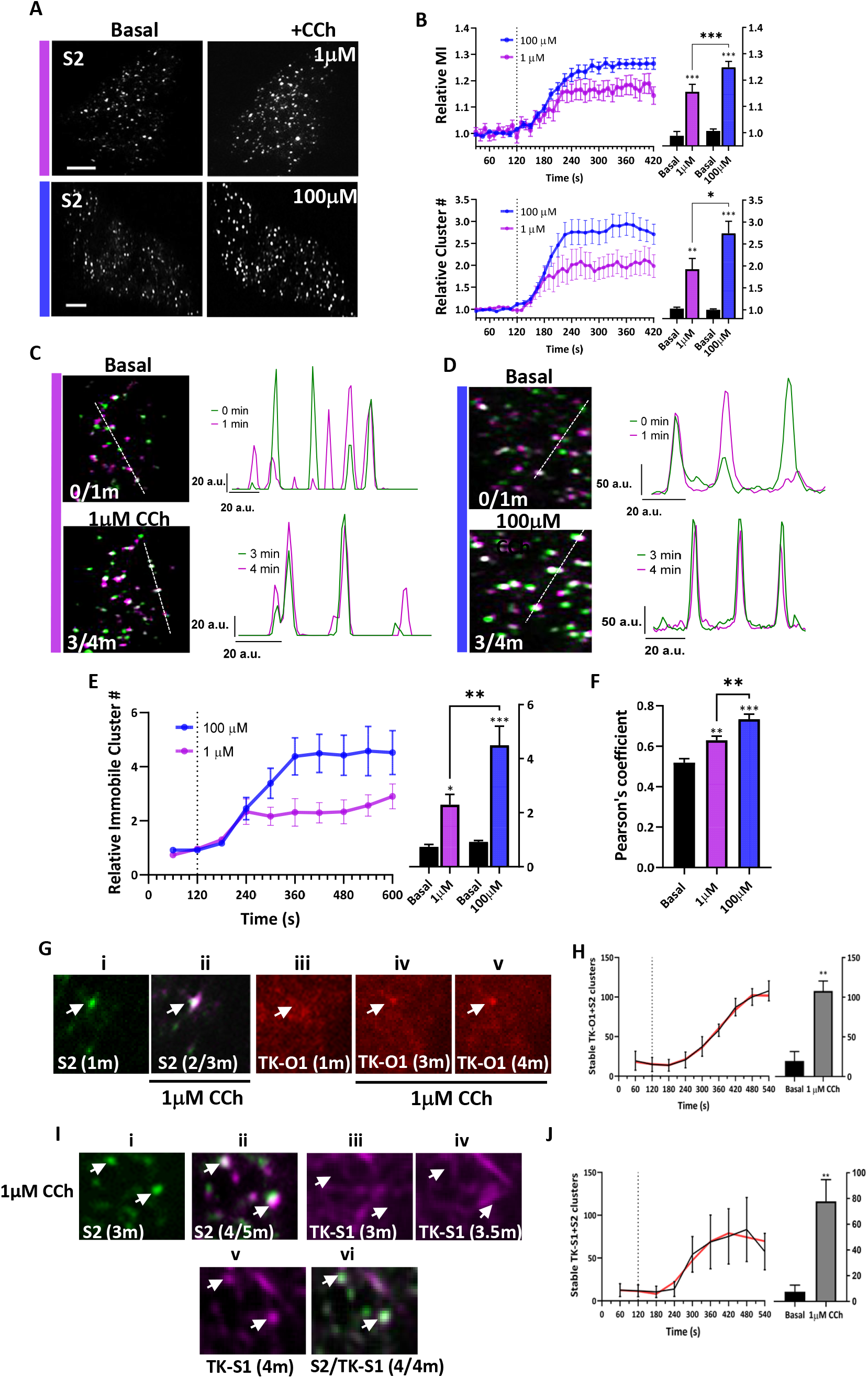
mVenus-STIM2 clusters display decreased mobility in response to carbachol stimulation. (*A*) mV-STIM2 in STIM2-KI cells (S2 in images) before (left) and after stimulation (with 1µM or 100µM CCh; right). (*B*) Time-dependent increase in mV-STIM2 cluster intensity (mean ± SEM of mean intensity, MI, upper panels) and number (Cluster #, lower panels) in response to 1µM (magenta) or 100µM CCh (blue) (CCh addition indicated by the dotted line). Respective bar graphs showing values at 1 min (basal) and 5 min (stimulated; 1µM or 100µM) time points. Scale bars, 5µm. (*C*) Overlay of mV-STIM2 images taken at 1 min intervals (clusters pseudocolored either magenta and green) with immobile clusters appearing as white. The overlay images are of frames before stimulation (0 and 1 min time points, 0/1m; upper) and after stimulation with 1µM CCh (3 and 4 min time points, 3/4m). Line scans on the right (position indicated in overlay images). (*D*) Images were analyzed as described in (*C*) for cells before and after stimulation with 100µM CCh. (*E*) Increase of immobile mV-STIM2 (mean ± SEM) clusters stimulated with 1µM (Magenta) or 100µM CCh (Blue). Bar graphs (right) show the number of clusters before and after stimulation. (*F*) Pearson correlation coefficients of clusters in (*C*) and (*D*), basal and after stimulation (1 and 5 min). (*G*) TK-O1 recruitment to immobile S2 clusters. S2 cluster (i) prior to stimulation (1 min) and (ii) overlay of S2 after stimulation with 1µM CCh (2/3 min overlay, immobile cluster indicated by arrow). TK-O1 cluster (iii) prior to stimulation and following stimulation at the (iv) 3 and (v) 4 min time points, with the emergent cluster co-localized with the immobile mV-STIM2 cluster (arrow). (*H*) Increase in immobile clusters containing both TK-O1 and S2 after 1µM CCh addition, bargraph showing number of immobile clusters before (basal) and after CCh stimulation (1 and 9 min time points, respectively, smoothened curve in Red). (*I*) TK-S1 recruitment to immobile S2 clusters. S2 clusters at (i) 3 min; (ii) overlay of images from 4 and 5 min (4/5m overlay) time points. TK-S1 clusters after CCh stimulation at (iii) 3, (iv) 3.5, and (v) 4 min time points. (vi) Overlay of TK-S1 and S2 at 4 min time point (4/4m). (*J*) Increase in immobile clusters containing both TK-O1 and S2 after 1µM CCh stimulation, bargraph showing number of immobile clusters before (basal) and after CCh stimulation (1 and 9 min time points respectively, smoothened curve (Red)). All TIRFM images show representative cells from at least 3 experiments with CCh added at 2 min. Statistical significance was assessed using Students t-tests for two groups and ANOVA for multiple groups, and presented as \**P* < 0.05, \*\**P* < 0.01, and \*\*\**P* < 0.001. Scale bars, 5µm.

The response of Orai1 to agonist stimulation in STIM2-KI cells was determined by expressing the proteins using the thymidine kinase (TK) promoter, a relatively weak promoter, to reduce the amount of expressed protein (17). While TK-Orai1-CFP (TK-O1) displayed a diffuse localization at the TIRF plane both in HEK293 cells (Fig. S3*A*) and S2KI-HEK293 cells (Fig. S3*B*), a few channel clusters could be seen prior to stimulation in both sets of cells (basal condition image and Fig. S3*A* and *B*). These TK-O1 clusters displayed overlap with mV-STIM2 clusters in basal STIM2-KI cells (Fig. S3*B,* upper images) and increased after stimulation with 1µM CCh (Fig. S3*A*, upper right image) together with co-clustering of both proteins (Fig. S3*B*, lower images and line scan). Note that immobile TK-O1 clusters were detected in STIM2-KI cells prior to stimulation (Fig. S3*C* white clusters, tracking was done for a min, 0/1m as described for Fig. 2) which increased following CCh stimulation (Fig. S3*D* shows overlay of TK-O1 signal 1 and 2 min after stimulation, frames 3/4m). Fig. S3 *E* and *F*, show mV-STIM2 in the same cell. Fig.2*G* shows recruitment of Orai1 to immobile STIM2 clusters (in this case, STIM2 clusters that were not pre-clustered with TK-O1 were tracked). (i, ii) A single STIM2 cluster that remained at the same location for a min after 1μM CCh stimulation (white cluster in second image indicates that it is immobile). (iii) Orai1 cluster was not detected prior to stimulation but seen at the same location as the mV-STIM2 cluster by 1 min after stimulation, staying at that location for another min (iv and v). Increase in number of immobile clusters containing TK-O1 and mV-STIM2 after 1μM CCh stimulation (Fig. 2*H*). Orai1 clustering in resting and stimulated cells was abrogated by silencing STIM2 (Fig. S3*A*, lower images). TK-O1clusters were also seen in unstimulated WT cells with increase after stimulation with 1μM CCh (Fig. S3 *H*). It is likely that the relatively low abundance of expressed Orai1 in the present study allowed detection of Orai1 clusters in resting cells.

TK-mCh-STIM1 (TK-S1), expressed in STIM2-KI cells. displayed the characteristic tracking pattern in cells under basal condition (Fig. S4 *A* and *B*, images in upper panel). TK-S1 (red) was not clustered with mV-STIM2 clusters (green) in basal cells (Fig. S4*A*) but co-clustered 5 min after 1μM CCh stimulation. TK-S1 clustering at 1μM CCh (images in second panel), but not at 100μM CCh stimulation (images in lower panel), was dependent on endogenous STIM2 (Fig. S4*B*). In contrast, basal clustering of mV-STIM2 and the increase after 1μM CCh was not dependent on endogenous STIM1 (Fig. S4*C*). Further, some TK-S1 clusters were seen 2 min after CCh stimulation (Fig. S4*D*) with a few immobile TK-S1 clusters detected 1 and 2 min after stimulation (Fig. S4*E*, overlay of frames 3/4m, mV-STIM2 clusters shown in Fig. S4*F*) with an increase in immobile clusters by 4-5 minutes of stimulation (Fig. *S4G* overlay of frames 6/7m, mV-STIM2 shown in Fig. S4*F*). Fig. 2*I* shows that STIM2 clusters (i) that were identified as immobile (ii; overlap of clusters between 2 and 3 min after stimulation, 4/5m) marked the site where TK-S1 co-clustered after stimulation (indicated by white arrows). TK-S1 clusters were detected at these locations only 2 min after stimulation (v), but not at earlier time points (iii and iv show images at 1 and 1.5 min after stimulation). The final panel (vi) shows co-localization of immobile STIM2 clusters with overlapping with newly formed STIM1 cluster (marked by white arrows). Time-dependent increase in number of immobile clusters containing both mV-STIM2 and TK-S1 following stimulation of cells with 1μM CCh is shown in Fig. 2*J*.

### E-Syts2/3-dependent ER-PM junctions mark the site of endogenous STIM2 clustering

Extended synaptotagmins 2 and 3 (E-Syts2/3) serve as constitutive tethers linking ER with the PM (18). Simultaneous knockdown of E-Syts2/3 expression in STIM2-KI cells greatly reduced mV- STIM2 clusters in resting cells (Fig. 3*A*). In contrast, overexpression of E-Syt2 or E-Syt3 increased mV-STIM2 clusters in unstimulated cells (Fig. 3 *B* and *C*). Localization of the ER or constitutive clusters of MAPPER were not dependent on endogenous STIM2, but was attenuated by loss of E- Syts2/3 expression (Fig. 3 *D* and *E*, respectively; stable pre-formed junctions, indicated by immobile mCerulean-MAPPER clusters, was decreased by knockdown of E-Syts2/3 (Fig. *4F*).

**Fig. 3.**
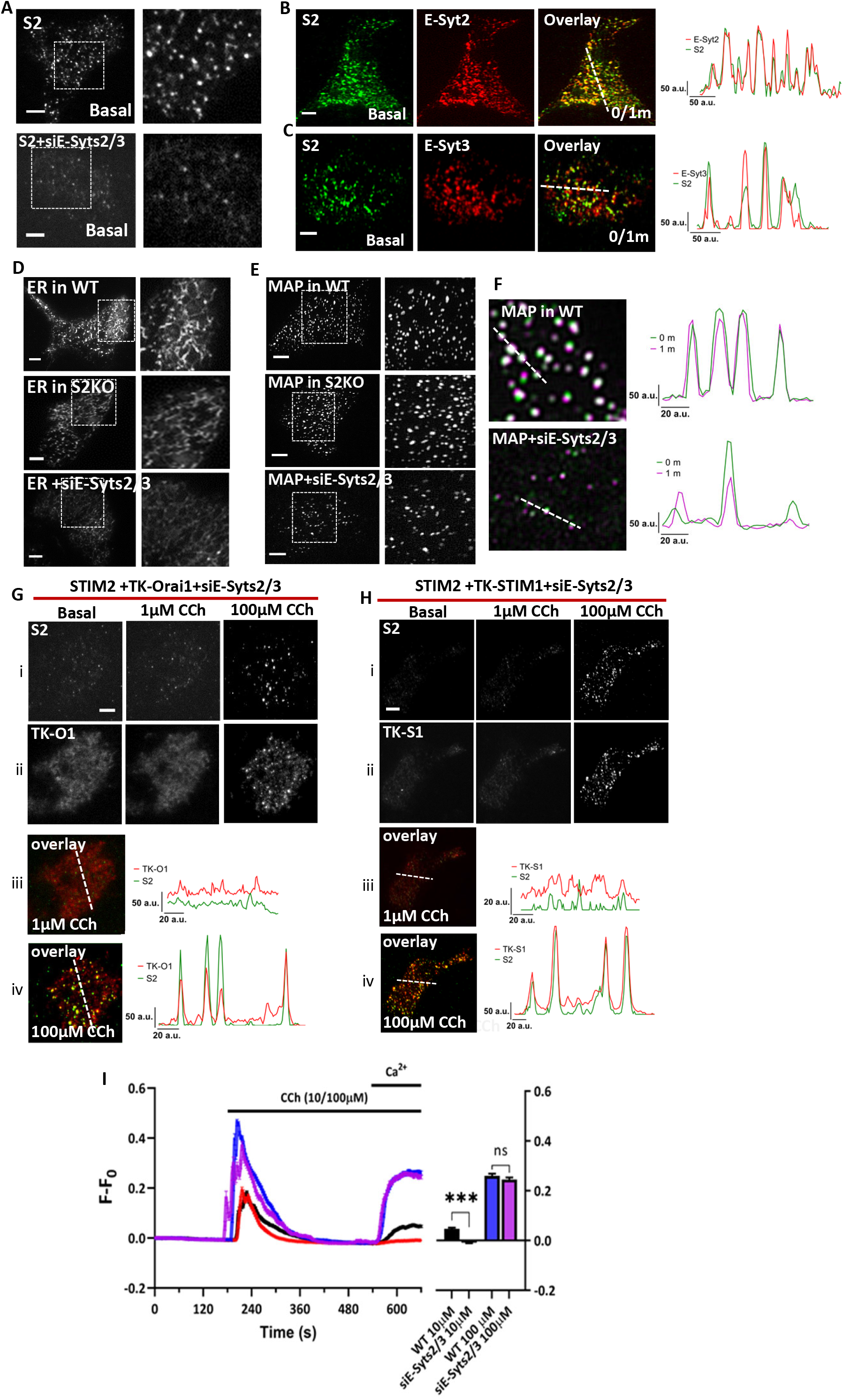
Role of extended synaptotagmins in localization of mVenus-STIM2 in ER-PM junction. (*A*) mV-STIM2 (S2) in control (top panel) and siE-Syts2/3-transfected STIM2-KI cells (lower panel); enlargement of demarcated area on right. (*B, C*) Unstimulated STIM2-KI cells expressing (*B*) mCh-E-Syt2 (ESyt2) or (*C*) mCh-E-Syt3 (ESyt2). From left to right: S2 (Green), ESyt2 or ESyt3 (Red), and overlay of both proteins showing shows colocalized clusters in yellow. Line scans (position indicated in overlay image) shows overlapping mV-S2 and E-Syt clusters. (*D*) mCherry-ER3 (ER) or (*E*) mCerulean-MAPPER (MAP) expressed in wild-type (WT, top panel) or STIM2 knockout (S2KO, middle panel) HEK293 cells, and WT cells treated with siE- Syts2/3 (bottom panel). Enlargements of the area demarcated by a square box are shown on the right. (*F*) Overlay images of mCerulean-MAPPER in WT cells (top) and WT cells treated with siE-Syts2/3 (bottom) under basal/unstimulated conditions (0/1m overlay) where white clusters represent immobile clusters. Line scans (position indicated in overlay image) show overlapping peaks for relatively immobile clusters. (*G*) STIM2-KI cells transfected with siE-Syts2/3 in addition to TK-Orai1-CFP; (i) mVenus-STIM2 (S2) and (ii) TK-Orai1-mCh (TK-O1) in unstimulated (left), 1µM CCh (middle) and 100µM conditions (right). (iii) and (iv) overlay of mV-STIM2 (S2; Green) and TK-Orai1-mCh (TK-O1; Red) following stimulation with 1µM and 100µM CCh, respectively. Line scans (position indicated on the overlay images) showing colocalized clusters of S2 and TK-O1. (*H*) Similar set of experiments as in (g) but with STIM2-KI cells expressing TK-mCh-STIM1 (TK-S1) + siE-Syts2/3. (*I*) Fura-2 fluorescence (F-F_0_, mean ± SEM) in control WT and siE-Syts2/3-treated WT cells stimulated with 10µM or 100µM CCh. Bar graphs shows increase in fluorescence due to 1mM Ca^2+^ entry. All TIRFM images show representative cells from at least 3 experiments, with CCh added at the 2 min time point. The Ca^2+^ imaging data are based on n >150 cells in each set. Statistical significance was assessed using Students t-tests for two groups and presented as not significant (n.s: *P* = 0.4) and significant (***: *P* < 0.001).

**Fig. 4.**
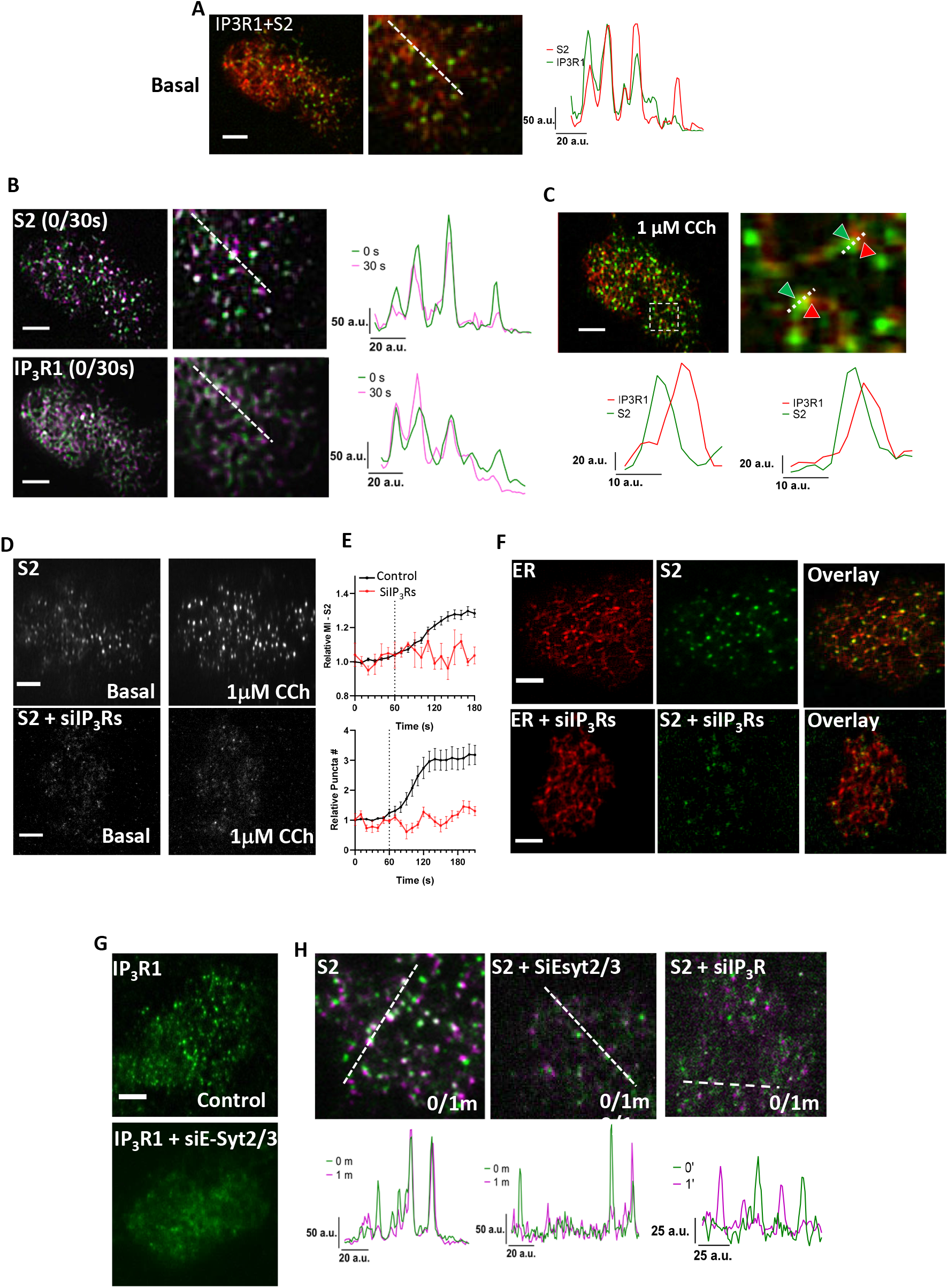
Clustering of STIM2 in the ER-PM junctional region depends on IP_3_Rs. (*A*) Overlay of mV-STIM2 (S2; Green) and IP_3_R1-mCherry (IP3R1; Red, expressed in STIM2-KI cells) with an enlarged image of demarcated area and line scan (position indicated on the enlarged overlay image). (*B*) Overlay images of S2 (top panel) and IP_3_R1-mCherry (IP_3_R1, lower panel) from 0 and 30s time points and line scans (position indicated on the enlarged overlay image). (*C*) Same cell as in *A* and *B* stimulated with 1μM CCh: overlay image of IP_3_R1 (Red) and S2 (Green) clusters and enlarged area (right). Line scan (below) show positions of the two proteins in two clusters (indicated by green arrow). (*D*) STIM2-KI cells showing S2 (upper panel) or cells with knockdown of all IP_3_R isoforms (lower panel). In each case, S2 fluorescence in unstimulated and 1µM CCh- stimulated cells are shown. (*E*) Increase in S2 cluster (mean intensity, top, and relative cluster number,bottom, in control (black trace) and IP_3_Rs knockdown cells (red trace)). (*F*) STIM2-KI cells expressing mCherry-ER3 showing S2 (Green) and mCherry-ER3 (ER, Red) in control (upper panels) and siIP_3_Rs-treated (lower panels) without stimulation. (*G*) Wild-type HEK293 transfected with IP_3_R1 or IP_3_R1+siE-Syts2/3. (*H*) Overlay images of S2 clusters in STIM2-KI cells alone, or with si-Esyt2/3 or siIP_3_Rs treatment from 0 and 1 min time points (0/1m). Line scan show overlapping peaks of both time points under basal condition. All TIRFM images show representative cells from at least 3 experiments. Scale bars, 5µm.

The increase in co-clustering of mV-STIM2 and TK-O1 following stimulation with 1μM CCh was substantially decreased in cells with knockdown of E-Syts2/3 (Fig. 4*G*, i, ii and iii; Supplementary Video 3) although there was some recovery of clustering following stimulation with 100μM CCh (Fig 4*G,* i, ii and iv). Similar effects of E-Syts2/3 knockdown were seen for co-clustering of TK- STIM1 and mV-STIM2 in response to stimulation with 1 and 100 µM CCh (Fig. 3*H*). Consistent with the effect of E-Syts2/3 knockdown on protein clustering, siE-Syts2/3 reduced Ca^2+^ entry stimulated by low but not high [CCh] (Fig. 3*I,* Ca^2+^ release was not affected). Together these data suggest that E-Syts2/3-mediated ER-PM tethering is required for STIM2 pre-clustering in ER-PM junctional region in basal conditions and following low intensity stimulation. Other tethering proteins, e.g. E-Syt1, might have a greater contribution to ER-PM junction assembly and STIM/Orai1 clustering at high stimulus intensities(18–20).

### Mobility of STIM2 clusters is dependent on IP_3_R

To examine whether other cellular cues, such as decrease in local [Ca^2+^]_ER_, are involved in constitutive STIM2 clustering, IP_3_R1-mCherry was expressed in STIM2-KI cells. Both proteins clustered in the ER-PM junctional region of unstimulated cells (Fig. 4*A*, line scans), with partial overlap between their clusters. We identified immobile mV-STIM2 clusters (Fig. *4B*, upper images and line scan) and immobile IP_3_R1 clusters (Fig. 4*B*, lower images and line scan) which appeared to be at the same cellular location. Magnified image of the clusters show that STIM2 clusters only partially overlapped with IP_3_R1 clusters (Fig.4*C*, note that images in Fig. 4 *A-C* were acquired from the same cell). Importantly, constitutive clustering of mV-STIM2 was strongly attenuated by knockdown of IP_3_Rs in unstimulated cells as well after 1μM CCh stimulation (Fig. 4*D*, Supplementary Video 4). Treatment with siRNAs against all three IP_3_R isoforms, siIP_3_Rs, also suppressed the time-dependent increase in mV-STIM2 clusters induced by in 1μM CCh stimulation (Fig. 4*E*). The presence of ER (indicated by mCh-ER3 signal in Fig. 4*F*) in the TIRF plane was not affected by knockdown of IP_3_Rs. Knockdown of E-Syts2/3, which disrupted localization of ER in the sub-plasma membrane region abrogated the clustering of IP_3_R1-mCherry (Fig. 4*G*) and reduced immobile clusters of STIM2 and IP_3_R1 in unstimulated cells (Fig. 4 *H* and *I*, respectively). These data reveal that pre-clustering of endogenous STIM2 in ER-PM junctions is dependent on IP_3_Rs and that ER-PM scaffolding by E-Syts2/3 by itself is not sufficient.

### Mobility of STIM2 clusters is determined by IP_3_R function and decrease of local [Ca^2+^]_ER_

Co-immunoprecipitation (co-IP) experiments revealed that neither Myc-tagged STIM1 nor STIM2 were pulled down together with IP_3_R1 (Fig. S5*A*, also shown by reverse co-IP in Fig. S5*B*). To assess whether IP_3_R function is involved in constitutive clustering of endogenous STIM2, STIM2- KI cells were treated with forskolin (FSK) which allosterically enhances IP_3_R channel actvity without an increase in [IP_3_] or IP_3_ binding by inducing PKA-dependent phosphorylation of IP_3_R1 and IP_3_R2 (21, 22). FSK (5µM) addition to STIM2-KI cells increased the number and mean fluorescence intensity of mV-STIM2 clusters (Fig. 5, upper two images in *A* and graph in *B;* Supplementary Video 5) and the number of immobile mV-STIM2 clusters (lower images in Fig. 5*A* show overlap of images acquired prior to (basal, 0/1m) and after FSK addition (3/4m)). Fig. 5*C* shows time-dependent FSK-induced increase in the number of mV-STIM2 clusters which was also detected when STIM2 (driven by TK promoter) was expressed in WT HEK-293 cells (Fig. 5*D*). Further, constitutive and FSK-induced STIM2 clustering were attenuated in cells lacking all three IP_3_Rs (IP_3_R-TKO) that were treated with siE-Syts2/3, although CPA-treatment restored STIM2 clustering (Fig. 5*E*, also see bar graphs).

**Fig. 5.**
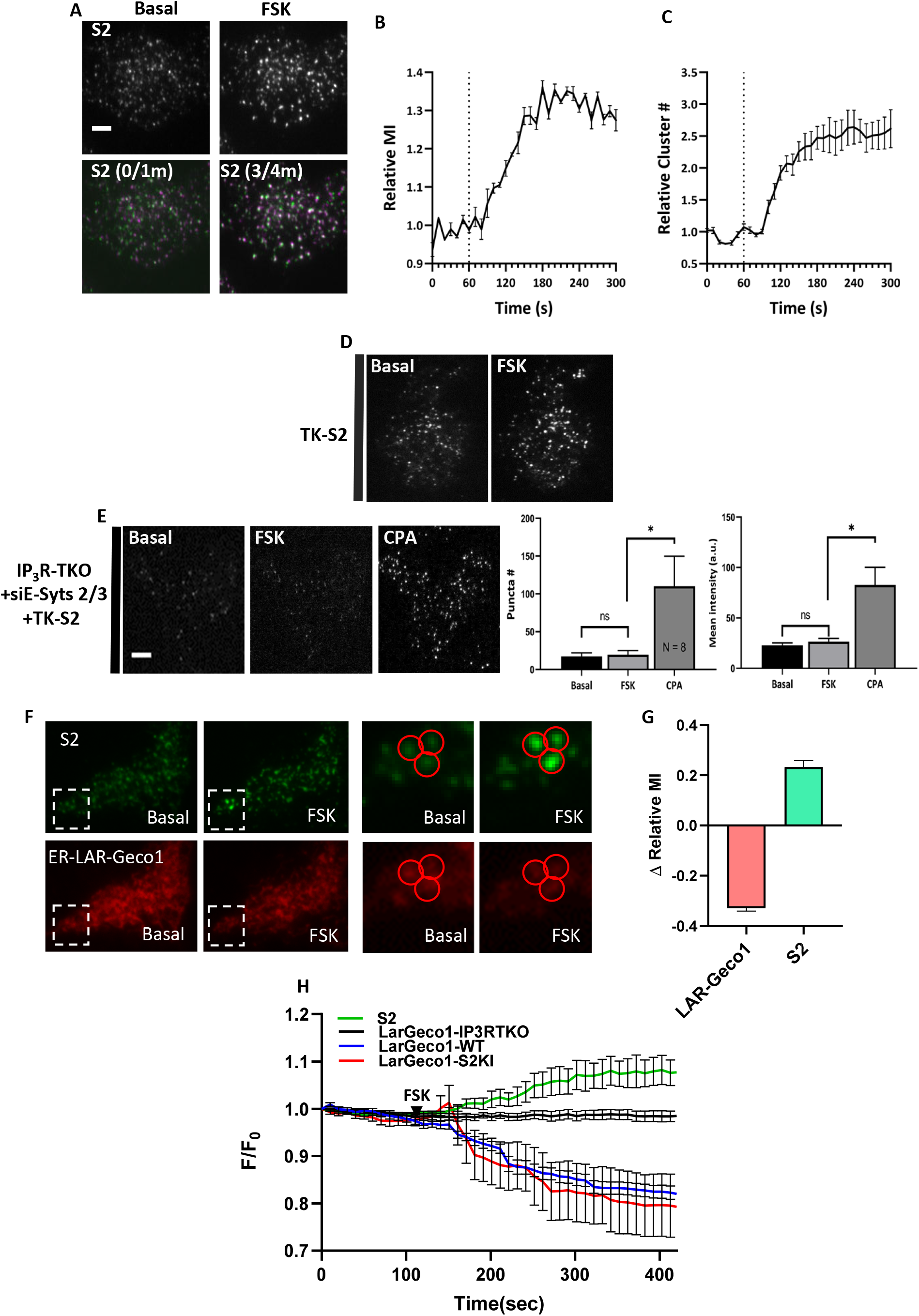
STIM2 clustering is determined by IP_3_R function. (*A*) mV-STIM2 clusters before and after 5µM forskolin (FSK) stimulation (top panel). Ooverlay images of S1 at 0 and 1 min (0/1m) and 3 and 4 min (3/4m) time points (bottom panel). (*B*) Increase of mV-S2 mean fluorescence intensity (MI) and relative number (Cluster #) (*C*) in response to FSK treatment (added at 1 min time point, dotted line). (*D*) HEK293 cells expressing TK-YFP-STIM2 (TK-S2) before and after stimulation with FSK. (*E*) IP_3_R-TKO cells (lack all three IP_3_R sub-types), treated with siE-Syts2/3 and expressing TK-S2, before and after stimulation with 5µM FSK and 25µM cyclopiazonic acid (CPA). Bar graphs showing number (Puncta #) and mean intensity intensity of TK-S2 clusters in the three conditions shown in the images (*n* =8). Statistical tests were done using ANOVA with the significance presented as not significant (n.s.; *P* > 0.05) and significant (\**P* < 0.05). (*F*) STIM2- KI cells expressing ER-Lar-GECO1: S2 (upper panel) and ER-Lar-GECO1 (lower panel) under basal (unstimulated) and stimulated with FSK (5µM). Enlargements of the region marked by a square are shown for visible camparisions. (*G*) Bar graph shows change in relative mean fluorescence intensity (MI) at the 5 min time point compared to basal (time point 1 min). Data is from 3 experiments and n=43 immobile clusters). (*H*) Line graphs showing whole cell intensity of ER- Lar-GECO1 in wild-type (WT), STIM2-KI (S2KI) and IP_3_R-TKO cells, and S2 fluorescence only in S2KI expressing Lar-GECO1. Addition of 5µM FSK is indicated by the black arrows. All TIRFM images show representative cells from at least 3 experiments. Scale bars, 5µm.

ER-Lar-GECO1 was used to measure changes in [Ca^2+^]_ER_ at the TIRF plane of STIM2-KI cells. Fig. 5*F* shows images of mV-STIM2 (upper set) and ER-Lar-GECO (lower set) in basal and FSK-treated STIM2-KI cells. The boxed highlights immobile mV-STIM2 clusters. ROI measurements of each immobile STIM2 puncta showed an increase in mV-STIM2 intensity and a decrease in ER-Lar-GECO1 intensity in response to FSK treatment (enlarged images in Fig. 5*F* and graph in Fig. 5*G*). Time course of this decrease following FSK addition to cells is shown in Fig. 5*H*. As a control experiment, we examined the change in ER-Lar-GECO1 intensity within the TIRF plane in IP_3_R-TKO cells, which did not display any response to FSK addition. The decrease in ER-Lar-GECO1 fluorescence seen in FSK-treated WT cells was comparable to that in STIM2-KI cells (Fig. 5*H* and Fig. S5 *C-E*).

### Determinants of IP_3_R function and STIM2 pre-clustering in resting cells

The role of IP_3_Rs in STIM2 clustering was further studied by expressing TK-YFP-STIM2 construct (TK-S2) in WT HEK293 cells or IP_3_R-TKO (23), IP_3_R-TKO stably expressing WT-IP_3_R1 (IP_3_R1), a pore dead mutant of IP_3_R1 (IP_3_R1G2506R) (24), or a phosphorylation-deficient mutant of IP_3_R1 (IP_3_R1- S1588A/S1755A, Phos-Mutant) (25). In WT cells, TK-YFP-STIM2 displayed constitutive clusters prior to any form of stimulation which was increased by FSK addition and further increased by CPA (Fig. 6*A*, first column; Supplementary Video 6). In the IP_3_R-TKO cells, TK-YFP-STIM2 clustering under basal conditions was reduced and did not increase after FSK addition, although subsequent addition of 25µM CPA did induce clustering of STIM2 (Fig. 6*A*, second column; Supplementary Video 7). Constitutive clustering of TK-YFP-STIM2 was also reduced in IP_3_R-TKO cells expressing pore-deficient IP_3_R1/G2506R and Phos-Mutants of IP_3_R. Further, FSK did not increase STIM2 clustering while subsequent addition of CPA did (Fig. 6*A*, third and fourth column). Constitutive clustering of TK-YFP-STIM2 was restored, together with the FSK- enhancement of clusters in IP_3_R-TKO cells stably expressing wild-type IP_3_R1 (Fig. 6*A*, last column; Supplementary Video 8). The relative increase in TK-S2 clusters (number and mean fluorescence intensity) seen in WT cells after FSK addition was significantly attenuated in IP_3_R-TKO cells or cells expressing mutant IP_3_Rs but was rescued in cells expressing IP_3_R1 (Fig. 6*B*). Time-dependent changes in STIM2 clustering following FSK stimulation in the different sets of cells are shown in Fig. 6 *C* and *D*, while Fig. 6*E* shows the effect of IP_3_R function on STIM2 cluster size in basal conditions and after FSK stimulation.

**Fig. 6.**
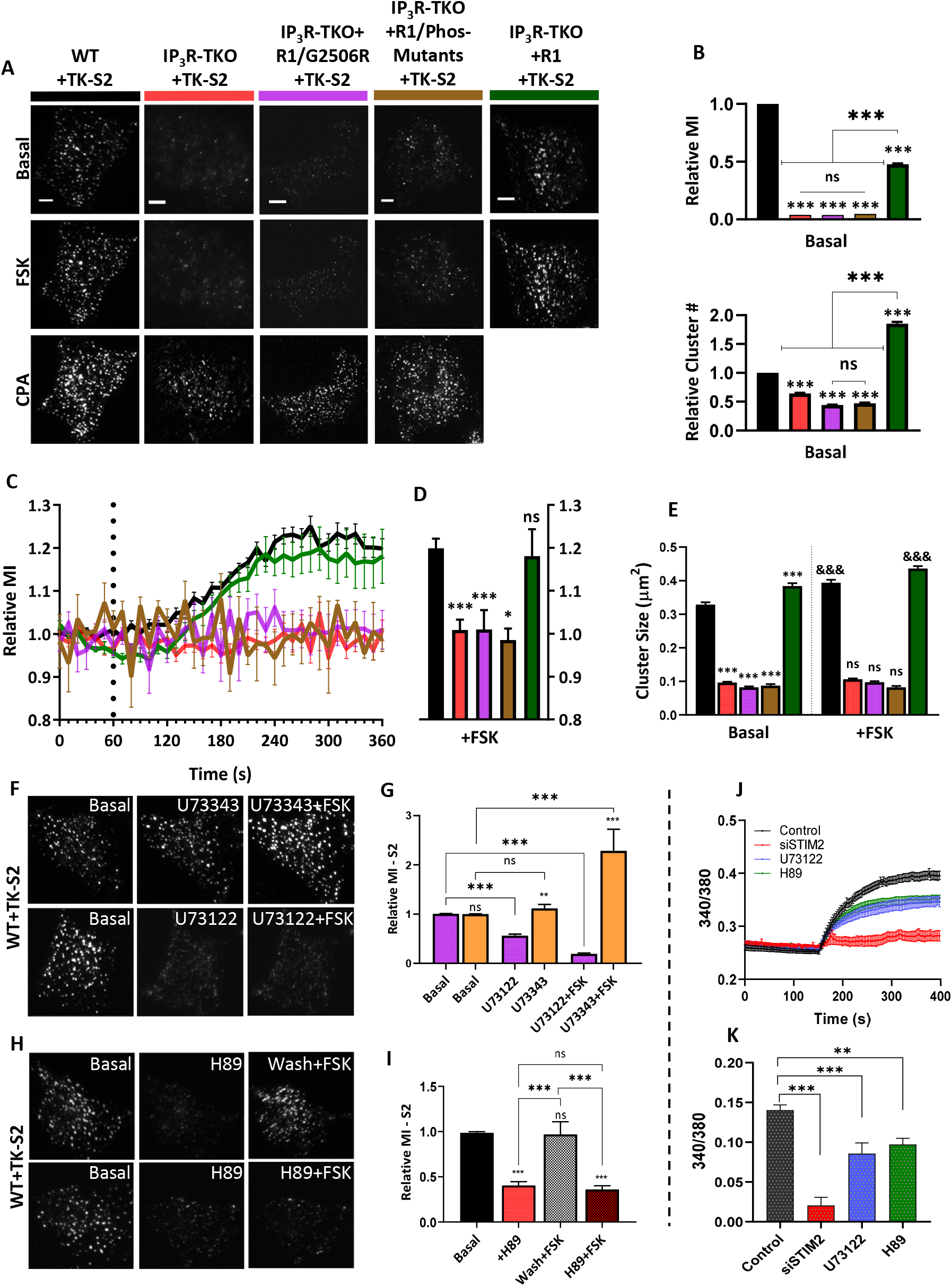
STIM2 clustering is determined by IP_3_R function. (*A*) HEK293 cells expressing TK- YFP-STIM2 (TK-S2): wild-type (WT, Black); IP_3_Rs-TKO (lack all three IP_3_R sub-types; Red); IP_3_R-TKO cells stably expressing a pore-dead mutant of IP_3_R1 (IP_3_R-TKO/G2506R, Magenta); IP_3_R-TKO cells expressing phosphorylation-deficient IP_3_R1 with S1588A and S1755A mutations (IP_3_R-TKO/Phos-Mutants, Brown); and IP_3_R-TKO cells stably expressing IP_3_R1 (IP_3_R- TKO+IP_3_R1, Green); respectively either unstimulated (basal) or consecutively stimulated with 5µM forskolin (FSK) and 25µM cyclopiazonic acid (CPA). (*B*) Bar graphs show relative mean intensity (MI; top) and relative number (bottom) of TK-S2 clusters in basal condition for all cell types in (*A*). (*C*) Time-dependent increases in relative MI of TK-S2 clusters calculated from cells shown in (*A*). (*D*) Bar graphs show relative MI at the 5 min time point in (*C*). (*E*) Bar graph showing relative size of TK-S2 clusters for the basal and FSK-treated cells in (*A*) Basal values for all IP_3_R-TKO cells were compared to WT, while FSK values for WT and all IP_3_R-TKO cells were compared with their basal counterparts. (*F*) WT cells expressing TK-S2 showing clusters at basal (left panel), and post-stimulation with either 5µM U73343 (inactive compound; middle panel) and U73343 with 5µM FSK (right panel). Lower set shows similar experiment done with the active PLC inhibitor, U73122 (5µM). (*G*) Bar graphs showing relative MI at basal and after stimulation with compounds as indicated in (*F*). (*H*) TK- S2 clusters in HEK293 cells under basal (left panel), post-stimulation with 5µM H89 (middle panel), and post-stimulation with 5µM FSK (right panel). Treatment with FSK was either with a washout (Wash) of H89 (top panel) or without (bottom panel). (*I*) Bar graphs showing relative MI at basal and after stimulation with compounds as indicated in (*H*). Statistical tests were done using ANOVA with non-significant results presented n.s. (*P* > 0.05) and various levels of statistical significance as &/* *P* < 0.05, \*\**P* < 0.01 and &&&/*** *P* < 0.001. All TIRFM images show representative cells from at least 3 experiments. Scale bars, 5µm. (*J*) Fura-2 fluorescence measurements presented as F-F_0_ (mean ± SEM) of S2KI (control, black) and S2KI+siSTIM2 cells (red) treated with 1mM CaCl (addition shown by arrow). Bar graphs show change in F-F_0_ due to 1mM Ca^2+^ entry (4 independent experiments with *n* > 170 cells).

The role of ambient PIP_2_-PLC activity in regulating IP_3_R function under basal conditions was assessed by treating WT HEK293 cells expressing TK-YFP-STIM2 with the PLC inhibitor, U73122. This compound, but not the inactive analogue, U73343, decreased STIM2 pre-clustering as well as FSK-induced enhancement of clustering (Fig. 6 *F* and *G*). These results were consistent with previous reports demonstrating PKA-dependent phosphorylation enhances IP_3_R function at low levels of IP_3_ (21). Treating cells with the PKA inhibitor, H89, suppressed the FSK-induced increase in TK-YFP-STIM2 clustering. These findings suggest that FSK-induced enhancement of STIM2 clustering is mediated via PKA-dependent phosphorylation of IP_3_R (see images and bar graphs in Fig. 6 *H* and *I*). Importantly, H89 alone also suppressed pre-clustering of STIM2 in the ER-PM junctional region. Consistent with these data, constitutive Ca^2+^ entry was significantly decreased by treatment with PLC and PKA inhibitors as well as by knockdown of endogenous STIM2 (Fig. 6 *J* and *K*). It is important to note that the contribution of cAMP/PKA to STIM2 pre-clustering in resting cells is likely to vary among different cell types. HEK293 cells have relatively high constitutive cAMP/PKA activity (26) and thus H89 was effective in reducing STIM2 pre-clustering in unstimulated cells. Other factors, such as the metabolic status (i.e. amount of ATP) that allosterically increase IP_3_R activity via enhancing the sensitivity of the receptor to IP_3_, could also act in a manner similar to PKA (27). We also cannot rule out a role for IP_3_R binding partners that enhance activity in the pre-clustering (28).

### STIM2-N terminus determines immobilization of STIM2 clusters in unstimulated cells

A STIM2 chimera where the N-terminus was replaced with that of STIM1 (S1N-S2C) and a STIM1 chimera containing STIM2 N-terminal domain (S2N-S1C) were used to determine whether STIM2 N-terminal EF-hand domain senses local [Ca^2+^]_ER_ calcium decrease under ambient conditions. TK-constructs of these chimeras were expressed in HEK cells. Although this STIM2 chimera (S1N-S2C) displayed pre-clustering in unstimulated cells, no immobile clusters were detected (Fig. 7*A*). In contrast, WT-STIM2 displayed robust pre-clustering with a population of immobile clusters (Fig *7B*, also see line-scans i and ii from the two sets of cells). Additionally, a STIM1 chimera with STIM2 N-terminus (S2N-S1C) displayed some pre-clusters with a few that were relatively immobile. Majority of S1-S2N appeared to be tracking as was the case for WT- STIM1. There were no detectable clusters in cells expressing WT-STIM1 (Fig. 7 *C* and *D*).

**Fig. 7.**
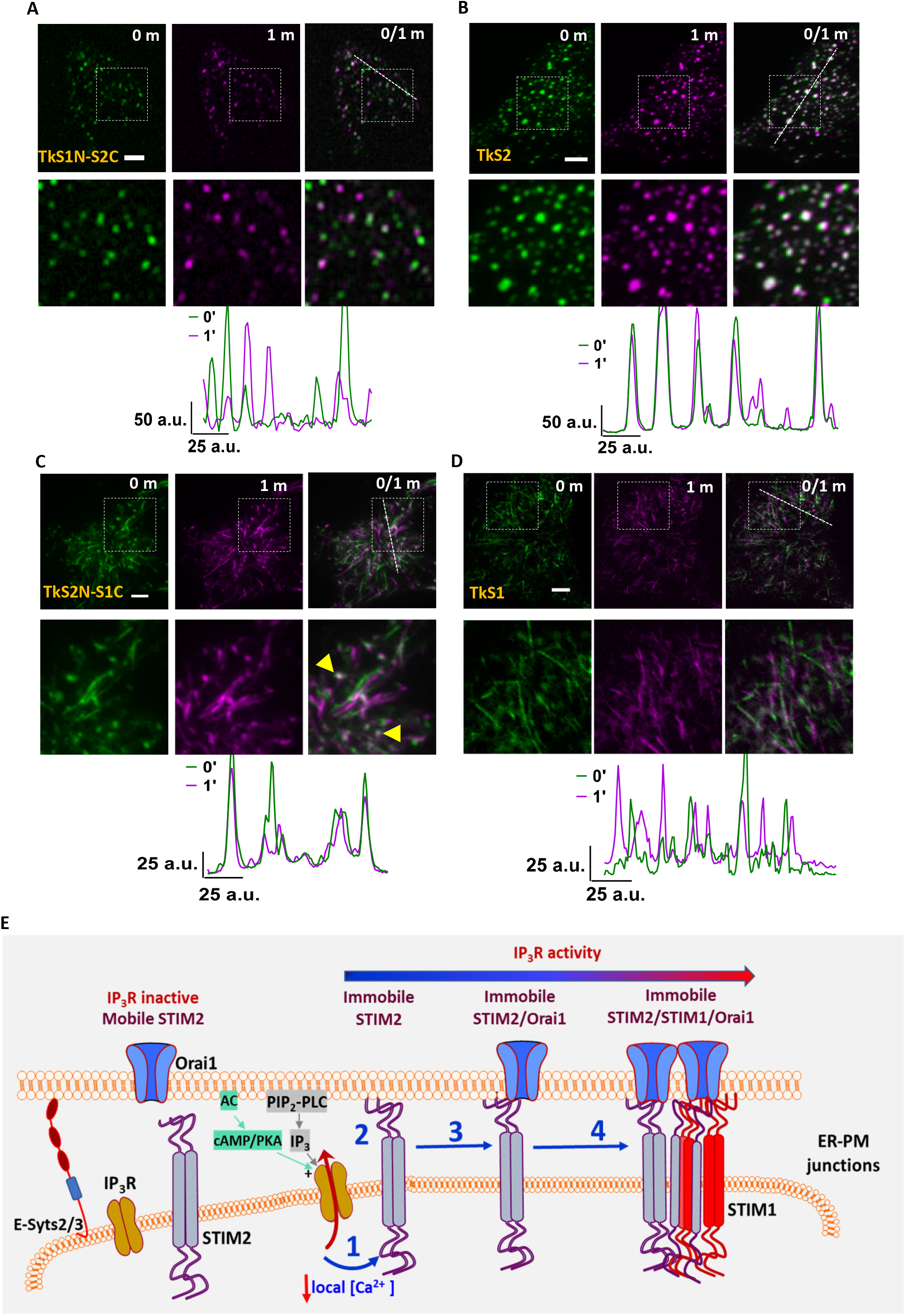
STIM2-N terminus senses local [Ca^2+^]_ER_ decreases at sub-threshold stimuli. HEK293 cells expressing (*A*) YFP-STIM2-STIM1N (STIM2 with STIM1 N-terminus, S2-S1N) or (*B*) YFP- STIM2 (Control, S2-WT) in basal conditions (0 and 1 min time points individually and with overlay (0/1m)). Enlargement of demarcated region shown in lower panel and line scans (position shown in enlarged images). HEK293 cells expressing (*C*) YFP-STIM1-STIM2N (STIM1 with STIM2- N-terminus, S1-S2N) or (*D*) YFP-STIM1 (Control, S1-WT) under basal conditions ( 0 and 1 min time points alone and with overlay (0/1m)). Enlargement of demarcated region shown in lower images and also line scans (position shown in enlarged images). All TIRFM images show representative cells from at least 3 experiments. Scale bars, 5µm. (*E*) Proposed Model: The illustration shows localization of IP_3_R and STIM2 in the ER-PM junctional region of the cell. Under ambient conditions (without agonist addition), constitutive PLC-dependent PIP_2_ hydrolysis as well as cAMP/PKA activity regulate the activation of IP_3_R. When STIM2 is in the vicinity of a functional IP_3_R, it senses the lower [Ca^2+^]_ER_ (1) which leads to scaffolding to the plasma membrane and immobilization (2). Orai1 is then recruited to immobile STIM2 (3) and with further [Ca^2+^]_ER_ decrease STIM1 is also recruited to immobile STIM2 (4). The model refers to cellular responses under ambient and low intensity stimuli. ER: endoplasmic reticulum; PM: plasma membrane.

## DISCUSSION

In the study described above, we utilized targeted gene editing to tag endogenous STIM2 with mVenus which enabled us to study of the distribution, dynamics, and regulation of the endogenous protein in unstimulated cells and after agonist stimulation. We show that endogenous STIM2 displays constitutive clustering within the sub-plasma membrane ER in resting cells under basal condition. Importantly, there is a small population of STIM2 clusters that is relatively immobile while majority of them are mobile. Our findings suggest that immobile STIM2 clusters are physiologically relevant as they are increased by agonist stimulation and contribute to initiation of SOCE in basal and agonist-stimulated cells. Further, localization of ER in the sub-plasma membrane region of cells is required for this clustering but it is not sufficient. Importantly, we have identified critical cellular clues which govern the clustering of STIM2 in cells under ambient conditions. A major finding of this study is that IP_3_R channel activity within sub-plasma membrane ER of cells underlies the constitutive clustering of STIM2. We show that immobile STIM2 clusters and immobile IP_3_R are juxtaposed and localized in sub-plasma membrane ER. As suggested previously, localization of functional immobile IP_3_R in sub-plasma membrane ER enables them to rapidly sense and respond to increases in [IP_3_] generated by hydrolysis of plasma membrane PIP_2_ (16). The close proximity of STIM2 and IP_3_R clusters allows the luminal N-terminus of STIM2 to sense IP_3_R-mediated decrease in local [Ca^2+^]_ER_, leading to immobilization of STIM2 in ER-PM junctions. Further, we also report that IP_3_R activity in unstimulated cells is maintained by IP_3_ that is generated by constitutive PLC-dependent PIP_2_ hydrolysis. Unexpectedly, PKA inhibitors also caused decrease in STIM2 pre-clustering, suggesting that cAMP/PKA signaling can also contribute to constitutive clustering of the protein. Consistent with this, addition of forskolin to cells increased the number of immobile STIM2 clusters in ER-PM junctional region.

Together, these important findings reveal that regulation of IP_3_R function under ambient conditions underlies constitutive clustering of STIM2. We also infer that IP_3_R activation under these conditions is likely to be transient, causing rapid local decrease in ER followed by refilling mediated by the SERCA pumps. In the absence of Ca^2+^ recycling in peripheral ER, e.g. in IP_3_R-TKO cells, STIM2 pre-clustering in ER-PM junctional region is suppressed. Similar suppression of immobile STIM2 clusters is seen when constitutive IP_3_R activity is reduced by inhibition of PLC or PKA or when Ca^2+^-sensitivity of STIM2 N-terminus is decreased. Conversely, preventing recycling by maximizing release enhances immobile clustering. Together, these findings reveal that continued recycling of Ca^2+^ in peripheral ER maintains constitutive clusters of STIM2 in the ER-PM junctional region, while sensing ER-Ca^2+^ store depletion causes immobilization of the protein.

We suggest that immobile STIM2 clusters represent the site of SOCE initiation in basal as well as agonist-stimulated conditions. These STIM2 clusters demarcate sites where Orai1 and STIM1 are recruited, which is especially relevant in cells under ambient conditions and following exposure to low intensity stimuli. Use of a weak promoter to drive Orai1 expression allowed detection of a few immobile Orai1 clusters that are co-localized with immobile mV-STIM2 clusters even in unstimulated cells and increased following agonist stimulation. Our data indicate that STIM2/Orai1 complex underlies basal Ca^2+^ entry in unstimulated cells since knockdown of endogenous STIM2 decreased basal Ca^2+^ entry as well as Orai1 clustering. In contrast, STIM1 clusters with immobile STIM2 only after agonist stimulation and prominent clustering is seen with longer stimulation times, i.e. with more depletion of [Ca^2+^]_ER_. Note that both Orai1 and STIM1 are stabilized within ER-PM junctions when co-clustered with the relatively immobile STIM2 and the number of stable complexes are increased in a dose- and time-dependent manner after agonist stimulation. Thus, immobilization of STIM2 clusters in response to local [Ca^2+^]_ER_ decrease marks a critical checkpoint for SOCE initiation. This change in STIM2 clustering is also the first step in coupling ER-Ca^2+^ store depletion with SOCE activation. Close association between STIM1 and IP_3_R has been shown in HEK293T cells, which enables a fast response of STIM1 to a decrease in [Ca^2+^]_ER_ following high-intensity stimuli (29). It is interesting to note that STIM2 forms complexes with RyR, STIM1 and Orai1 in T cells, which determine generation of Ca^2+^ microdomains following T cell receptor stimulation (30). Conversely, Ca^2+^ binding N-terminal region of STIM proteins is suggested to suppress IP_3_R function under resting conditions. This inhibition is relieved when STIMs are activated with increasing agonist concentrations (31). During the preparation of this manuscript, a mathematical model was described to explain the spontaneous Ca^2+^ microdomains that are detected in unstimulated T cells (32). According to this model, pre-formed clusters of STIM2/Orai1 are localized in close proximity to IP_3_Rs which facilitates STIM2 to detect reduction in local [Ca^2+^]_ER_ caused by spontaneous IP_3_R activity. However, no experimental data were provided to support this model. The findings we report herein provide evidence that IP_3_R activity in unstimulated cells is not spontaneous but rather that it is regulated by intracellular signals generated under ambient conditions. While further studies are required to determine the exact factors that govern localization of functional IP_3_R near ER-PM junctions, our data reveal that STIM2 serves as the crucial first responder that senses the decrease in local ER-[Ca^2+^] mediated by IP_3_Rs. This functional communication between IP_3_R and STIM2 enables cells to distinguish between noise and a bona fide signal for activation of SOCE.

In conclusion, we have demonstrated here that pre-clustering of STIM2 in ER-PM junctions of unstimulated cells is not a random event but rather it is orchestrated by IP_3_R function via decrease in local [Ca^2+^]_ER_ which is sensed by the STIM2 N-terminus. Further, IP_3_R activity, and therefore STIM2 pre-clustering, in cells in absence of agonist stimulation is controlled by ambient PLC-dependent PM PIP_2_ hydrolysis together with constitutive cAMP/PKA activity. As noted above, depending on the cell type, other factors that allosterically increase IP_3_R activity, can also act in a manner similar to PKA phosphorylation. We show that a small population of STIM2 clusters is immobilized within ER-PM junctions in response to decrease in local [Ca^2+^]_ER_ which likely represent the active state of STIM2. This is further illustrated in our model (Fig. 7*E*) which depicts STIM2 clusters localized in ER-PM junctional region together with IP_3_R. When a mobile STIM2 cluster is in the vicinity of an activated “licensed” IP_3_R, it senses the decrease in local [Ca^2+^]_ER_ and responds causing scaffolding of its C-terminal polybasic domain to PM PIP_2_. This results in immobilization of the STIM2 cluster. Both Orai1 and STIM1 converge on these immobile STIM2 clusters and once co-clustered with STIM2, these proteins also display decreased mobility. The relatively flexible C-terminus of STIM2 (10, 33), as well as the lower Ca^2+^ affinity of its N-terminal EF-hand domain (4), together with the juxtaposition of IP_3_R in the ER-PM junctional region, allows it to sense and rapidly respond to decreases in local [Ca^2+^]_ER_. This critical functional link between IP_3_R and STIM2 underlies the constitutive clustering of STIM2 and ensures the coupling of ER-Ca^2+^ store release events with activation of SOCE and regulation of Ca^2+^ signaling in the cell.

## MATERIALS AND METHODS

### Cell culture, plasmids, RNAi transfection and reagents

HEK293 cells were maintained and transfected with various siRNAS or plasmids as described in Sl section.

### Knock-in of m-Venus in STIM2 gene

The standard method for CRISPR/Cas9 based approach was followed to knock-in the m-Venus tag at STIM2-N terminus. Both 5’ and 3’ homology repair template were generated by PCR, synthesized mVenus fragments were assembled using HiFi Infusion Kit (New England Biolabs, Ipswich, MA) into linearized pUC19 vector (Takara Bio USA, Inc) and confirmed by sequencing. Other details of the method including that for cell sorting and establishing single cell clonal line are provided in SI.

### Cellular imaging and image analysis

[Ca^2+^]_i_ imaging measurements, confocal microscopy, ZEISS AiryScan, and TIRFM were all used for live-cell imaging experiments described in the text and figure legends (details of microscope, acquisition and image analysis are provided in SI) **Statistics.** Data analysis was performed using Origin (OriginLab Corporation) and GraphPad (GraphPad Software). Statistical comparisons between two groups were made using the Student’s t-test, whereas comparisons of multiple groups were made using ANOVA followed by Sidak multiple comparisons test. Statistical significance was shown as significant at \**P* < 0.05, \*\**P* < 0.01 and \*\*\**P* < 0.001, or non-significant at *P* > 0.05 (n.s.).

## Acknowledgements

We would like to thank the following people for kindly providing the constructs used in this study: Dr. Tamas Balla (NICHD, NIH, Bethesda, MD, USA) for Orai1-LL-CFP; Dr. Tobias Meyer (Stanford University) for YFP-STIM2; Dr. Richard Lewis (Stanford University, Stanford, CA, USA) for mCherry-STIM1; and Dr. Jen Liou (UT Southwestern, Dallas, TX) GFP-MAPPER for GFP-MAPPER. Funding for this work was provided by NIDCR-DIR, NIH (Z01-DE00438-33) for I.A.; NIH grants: R35-HL150778 to M.T. and R01 DE019245 to D.I.Y.

## Author Contributions

MA - generated knock-in cell line, planned experiments, carried out imaging of cells, analyzed images and data. prepared manuscript (text, figures etc.)

HLO – contributed to imaging, wrote the script for image analysis, analyzed images and data. prepared manuscript (text, figures etc.)

HS - analyzed data, prepared manuscript (text, figures etc.)

G-YS – carried out calcium experiments and analyzed data

ZS - carried out calcium experiments and analyzed data

LP – generated IP_3_RTKO and other cell lines and carried out Co-IP experiments with IP3Rs and STIM proteins.

MT – provided cell line, contributed to manuscript preparation and discussion

DIY - provided cell line and other reagents, planned experiments, contributed to data discussion and manuscript preparation

IA- planned and supervised all experimental work, data interpretation and analysis, wrote manuscript.

## Supplementary Information

### Supplemental Materials and Methods

#### Cell culture, plasmids, RNAi transfection and reagents

HEK293 cells were cultured in DMEM supplemented with 10% heat-inactivated fetal bovine serum, 1% glutamine, and 1% penicillin/streptomycin, incubation at 37°C under 10% CO_2._ Lipofectamine RNAiMAX (Life Technologies, Carlsbad, CA, USA) was used for siRNA transfections. Mock transfections of control cells were conducted using negative control siRNA (Invitrogen). Cells were typically used 48 h post-transfection before imaging. The mCherry-ER3, mCerulean-MAPPER, IP3R1-mCh, pore-dead (G2506R) and phosphorylation deficient (S1588A and S1755A; Phos-Mutant) mutant of IP_3_R1, TK-Orai1-CFP, TK-mCherry-STIM1, ER-Lar-GECO1, YFP-STIM2, YFP-STIM1, YFP-STIM1-S2N, YFP-STIM2-S1N or TK-YFP-STIM2 plasmids were transfected using electroporation following the protocol recommended by Lonza (Walkersville, MD, USA). Mouse monoclonal antibody of STIM1 and STIM2 antibodies were obtained from Cell Signaling Technology (Danvers, MA, USA). Rabbit polyclonal antibody to Orai1 was produced against C-terminal and purified by ProSci (Poway, CA, USA) and used described previously (1, 2). All other reagents used were of molecular biology grade obtained from Sigma-Aldrich (St Louis, MO, USA) unless mentioned otherwise.

#### CRISPR/Cas9 approach for mVenus Knock-in at C-terminus of STIM2 gene

The standard method for CRISPR/Cas9 based Knock-in approach was followed (3). Both 5’ and 3’ homology repair template were generated by PCR, synthesized mVenus fragments were assembled using HiFi Infusion Kit (New England Biolabs, Ipswich, MA) into linearized pUC19 vector (Takara Bio USA, Inc) and confirmed by sequencing. Multiple gRNA was designed by GPP sgRNA Designer of the Broad Institute and Benchling software (Benchling, San Francisco, USA). gRNA sequence was cloned in pX330 vector obtained from Addgene (Watertown, MA). Electroporation of HDR template and gRNA in and HEK293 cell line and fluorescence microscopy was performed after 2- 3 days. Fluorescent cell sorting was done to enrich the fluorescent population. Single-cell clones were obtained by sorting a single cell in each well of a 96 well plate. At least five different clones were further confirmed by PCR, sequencing, and endogenous protein tagged with mVenus expression was confirmed by western blot analysis with GFP antibody. A homozygous single cell population was used in this study.

#### Plasmids and Cell lines

The mCherry-ER3 plasmid was obtained from Addgene (Watertown, MA). To generate the TK promoter-driven constructs, the following plasmids were used as templates: Orai1-LL-CFP (Tamas Balla, NICHD, NIH, Bethesda, MD, USA), YFP-STIM2 (Tobias Meyer, Stanford University) and mCherry-STIM1 (Richard Lewis, Stanford University, Stanford, CA, USA). Briefly, the CMV promoter in these plasmids were substituted with the TK promoter. To generate mCerulean-MAPPER, the fluorescent tag in GFP-MAPPER (Jen Liou, UT Southwestern, Dallas, TX) was substituted with mCerulean (Addgene). These constructs were generated and then sequenced to confirm proper substitutions of the promoter or fluorescent tag by Mutagenex Corporation (Suwanee, GA). mCh-IP3R1, as well as IP3R-TKO (Null), IP3R-TKO/hIP3R1, IP3R1-TKO/R1G2506R, IP3R-TKO/IP3R1-Phos-Mutants are characterized previously (4–6).

#### [Ca^2+^]_i_ measurements

Fura*-2* fluorescence was measured in cells cultured for 24 h in collagen-coated glass-bottomed MatTek tissue culture dishes (MatTek Corp. Ashland, MA, USA) and transfected as required. All experiments were done 24–48 h post-transfection. Wells were loaded with 1µM Fura-2AM (Life Technologies) for 30 min at 37 °C. Fluorescence was measured with the Olympus IX50 microscope (Olympus, Center Valley, PA), Polychrome V (Till Photonics LLC, Pleasanton, CA), and EM-CCD camera (Hamamatsu, Tokyo, Japan). Image acquisition and data processing were done using MetaFlour (Molecular Devices, Downingtown, PA). Additions of carbachol (1, 10 or 100µM) and 1mM CaCl_2_ are indicated by arrows on the graphs.

#### Confocal Microscopy

The FV3000 confocal laser scanning microscope (Olympus) was used to acquire live cell images of the STIM2-KI cell line (expressing mV-STIM2). All cells were imaged at 37°C using a Plan-Apochromat 63×1.4 NA oil-immersion objective lens. The imaging software, FluoView, was used to excite mVenus using the 514nm laser and collect the subsequent emitted light. Images were further analyzed using Fiji/ImageJ2.

#### ZEISS AiryScan Imaging

Airyscan imaging was performed with LSM880, a confocal laser scanning microscope ZEISS LSM 880 (Carl Zeiss AG, Oberkochen, Germany) equipped with an Airyscan detection unit. Airyscan technology is an improved high-resolution imaging, which can acquire images with up to 8x improvement in Signal to Noise ratio and 1.7x higher resolution than a conventional confocal. To maximize the resolution enhancement, 63X/1.46 Oil Corr M27 or Plan-Apochromat 63X/1.40 Oil Corr M27 objectives (Zeiss) with Immersol 518 F immersion media (n_e_ = 1.518 (23 °C); Carl Zeiss) was used for the imaging.

#### TIRF Microscopy (TIRFM) Imaging

TIRF microscopy experiments were using HEK293 cells cultured for 24h on glass-bottomed dishes coated with collagen (MatTek) and transfected/electroporated as required. All experiments were done 24-48h post-transfection. TIRFM was performed using an Olympus IX81 motorized inverted microscope (Olympus) with a TIRF-optimized Olympus Plan APO 60× (1.45 NA) oil immersion objective and Lambda 10-3 filter wheel (Sutter Instruments, Novato, CA, USA) containing 480-band pass (BP 40 m), 540- band pass (BP 30 m), and 575lp emission filters (Chroma Technology, Bellows Falls, VT). Images were collected using a Hamamatsu ORCA-Flash4.0 camera (Olympus) and the MetaMorph imaging software (Molecular Devices), which were then analyzed as described below.

#### Image Analysis

The Zen Black 2.1 software (Carl Zeiss) was used to acquire and process images on the Airyscan microscope imaging system. Data acquired by each of the 32 Airy detector channels were processed separately by performing filtering, deconvolution and pixel reassignment in order to obtain images with enhanced SNR and improved spatial resolution. For TIRFM imaging, both MetaMorph and Fiji/ImageJ2 were used in the analysis. For STIM1 and STIM2, the image stacks were first processed in MetaMorph using a region-of-interest (ROI)-based background subtraction method. For Orai1, the image stack was first processed using a Gaussian Blur filter (sigma = 5) and then used to generate an average intensity projection of the entire stack, which was then subtracted from each image in the stack. All background-corrected image stacks were analyzed in Fiji/ImageJ2 using macros developed in-house to obtain two measurement parameters: the number of clusters detected and mean fluorescence intensity. The image stack was run through the FeatureJ/Laplacian filter (7) and auto-thresholded using the Otsu method to generate a binary image stack where each cluster was displayed as white particles on a black background. Using the Set Measurements and Analyze Particles commands, ROIs were drawn around each white particle and saved as a ROI set specific for each binary image. The ROI sets were then applied to the background-corrected image to measure mean fluorescence intensity. The STIM2 images were used to generate these ROI sets, which were then applied to other images of STIM1 and/or Orai1 if these proteins were also co-expressed in the same cell. The number of ROIs generated for each image was calculated as number of clusters detected. The quantitative measurements for each image were saved as a comma separated value-formatted files, which were then imported into Origin (OriginLab Corporation, Northampton, MA, USA) and GraphPad Prism (GraphPad Software, La Jolla, CA, USA) for further analyses. Whole Cell ROI was also performed, wherever indicated in result section, to quantify the whole cell fluorescence intensity from basal to stimulated cells.

Two approaches were used to identify clusters that are stable for at least 1 min. Two frames of 1 min interval were pseudo-colored green and magenta to create overlay images. Mobile clusters will be either green or magenta, while immobile clusters will be white. Numeric time values (in min, labeled as m) stated on the images are counted up from the start of imaging. For example, ‘0/1m’ represent images from the start of the experiment (0 min) and a min later (1 min). Line scans were performed in Fiji/ImageJ2 by drawing a line across the image and importing the underlying plot data into Graphpad Prism for graphing. The Image Calculator tool in Fiji was used to apply a logical AND on consecutive binary images (at 1 min intervals) to identify objects having the same position in both images. The Set Measurements and Analyze Particles commands were then used as described above to calculate the number of immobile clusters detected. Overlay images were analyzed using Coloc2 plugin in Fiji/ImageJ2 to get Pearson’s Correlation Coefficient (PCC) and Mander’s Overlap Coefficient (MOC) values.

#### Co-immunoprecipitation (Co-IP) and Western blotting

Wild-type (WT) and STIM2-KI HEK293 cells were treated as indicated in the legend, washed with 1× phosphate-buffered saline and lysed in Pierce IP lysis buffer supplemented with protease inhibitors (all materials were from Thermo Fisher Scientific, Waltham, MA). Cell lysates were centrifuged (10,000 g, 10 min at 4 °C) and quantified by BCA protein assay kit (Thermo Scientific). Co-IP experiments were done using anti-Myc magnetic beads, or Protein A Sepharose CL-4B (GE Healthcare Life Sciences, Marlborough, MA) as described earlier unless indicated otherwise. For Myc-tagged proteins, Pierce anti-Myc magnetic beads (Thermo Scientific) were used following the manufacturer’s instruction. After wash the immunoprecipitants were eluted in elution buffer or in gel-loading buffer by heating at 95 °C for 5-10 min and resolved in 4–12% NuPAGE gels (Life Technologies), followed by Western blotting. Proteins of interest were immunoblotted using anti-IP_3_R1 and anti-Myc, anti-STIM2 (Cell Signaling Technologies, Danvers, MA), anti-β-actin (Abcam, Cambridge, MA) antibodies.

#### Statistics

Data analysis was performed using Origin (OriginLab Corporation) and GraphPad (GraphPad Software). Relative values have been expressed in comparison to the basal condition (average value of time frame before stimulation in timelapse analysis) or to the WT counterpart (in static analysis). Smoothened curves were generated using GraphPad to help visualize trends where appropriate and were not used for any statistical analysis. Experimental values are expressed as mean ± SEM. Statistical comparisons between two groups were made using the Student’s t-test, whereas comparisons of multiple groups were made using one-way ANOVA followed by Sidak multiple comparisons test. Statistical significance was shown as significant at \**P* < 0.05, \*\**P* < 0.01 and \*\*\**P* < 0.001, or non-significant at *P* > 0.05 (n.s.).

### Supplementary Figure Legends

**Fig. S1.**
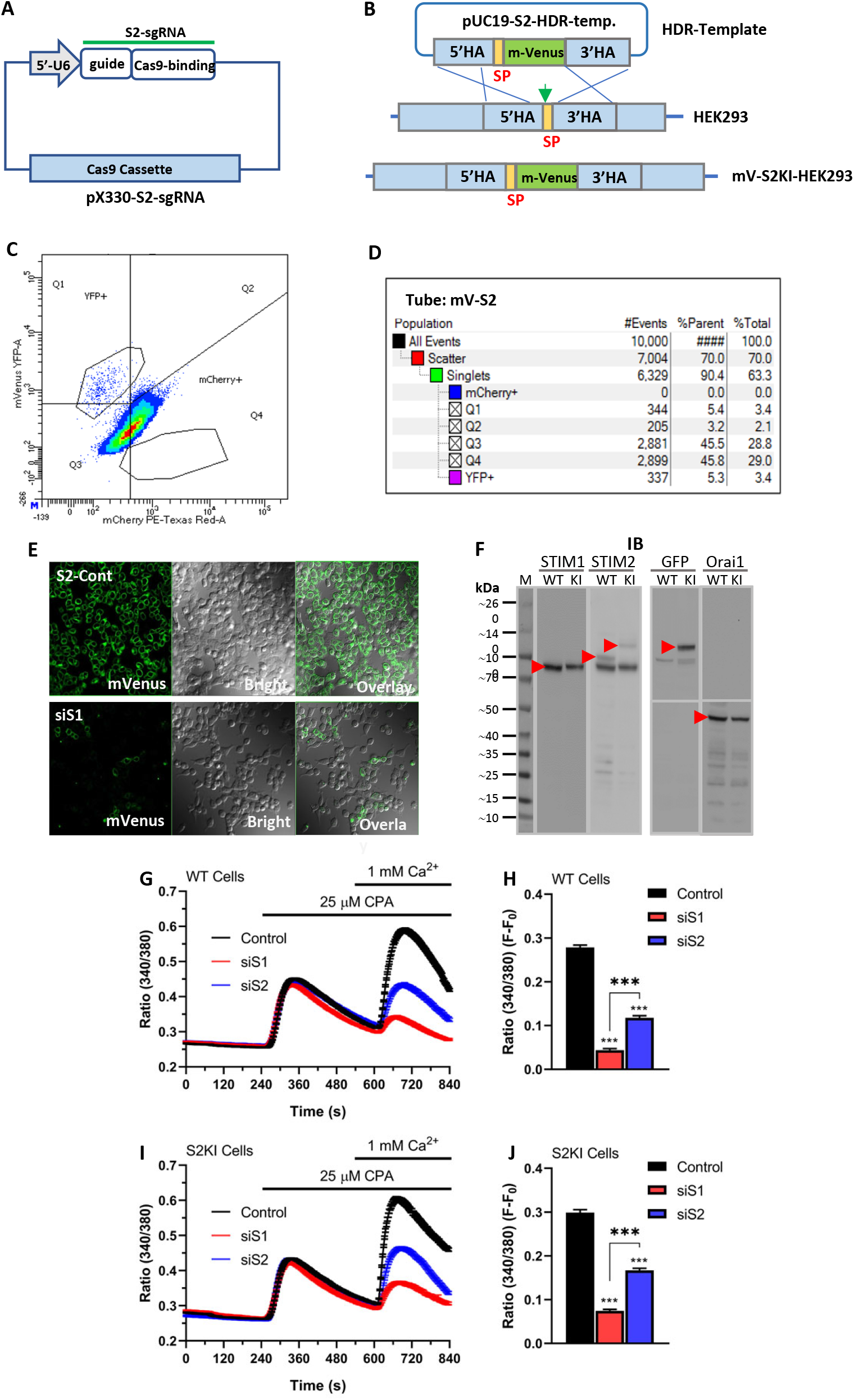
Generation and validation of mVenus-STIM2 knock-in cell line (STIM2-KI) using the CRISPR/Cas9 strategy. (*A*) Schematic of plasmid construct showing gRNA and cas9 expression cassette. (*B*) Schematic of donor plasmid as repair template for knock-in of mVenus immediately after the signal peptide of STIM2. Site-specific cleavage of *Stim2* by the Cas9-sgRNA complex in wild-type (WT) HEK293 cells is repaired through homology-directed repair with homology arms that flank an in frame mVenus open reading frame. (*C, D*) Flow cytometry-based sorting of mVenus-STIM2 v(mV-STIM2)-positive cells (STIM2-KI cells) and subsequently single cell cloning using a 96-well plate to generate a monoclonal population. (*E*) mVenus-STIM2 expression; in STIM2-KI without (S2-Cont) and with siSTIM2 treatment (siS2). The images are representative of data obtained from 3 experiments. (*F*) Western blots showing expression of STIM2, mV-STIM2 (endogenous STIM2), STIM1, and Orai1 in wild-type (WT) and STIM2-KI (KI) cells using anti-STIM1, anti-STIM2 (same blot used after detection of STIM1), anti-GFP (also detects mVenus-STIM2) and anti-Orai1 antibodies. Red arrows indicate location of the respective proteins on the blots. (*G-J*) Fura-2 fluorescence measurements using the Ca^2+^ addback assay where cells were stimulated with cyclopiazonic acid (CPA; 25µM) in Ca^2+^-free medium and then with 1 mM CaCl_2_ (Ca^2+^) in (*G, H*) WT HEK293 cells and (*I, J*) STIM2-KI (S2-KI) cells. All additions are as indicated by the black arrows. Black traces are control cells, while red and blue traces indicate cells treated with siSTIM1 and siSTIM2 respectively. (*H, J*) Bar graphs showing calcium entry components (F-F_0_) in the cells shown in *G* and *H*. Statistical significance was assessed using ANOVA for multiple groups and presented as not significant (n.s.; *P* > 0.05) and significant (\*\*\**P* < 0.001). Graphs show averaged data (mean ± SEM) obtained from n >300 cells from at least 3 experiments.

**Fig. S2.**
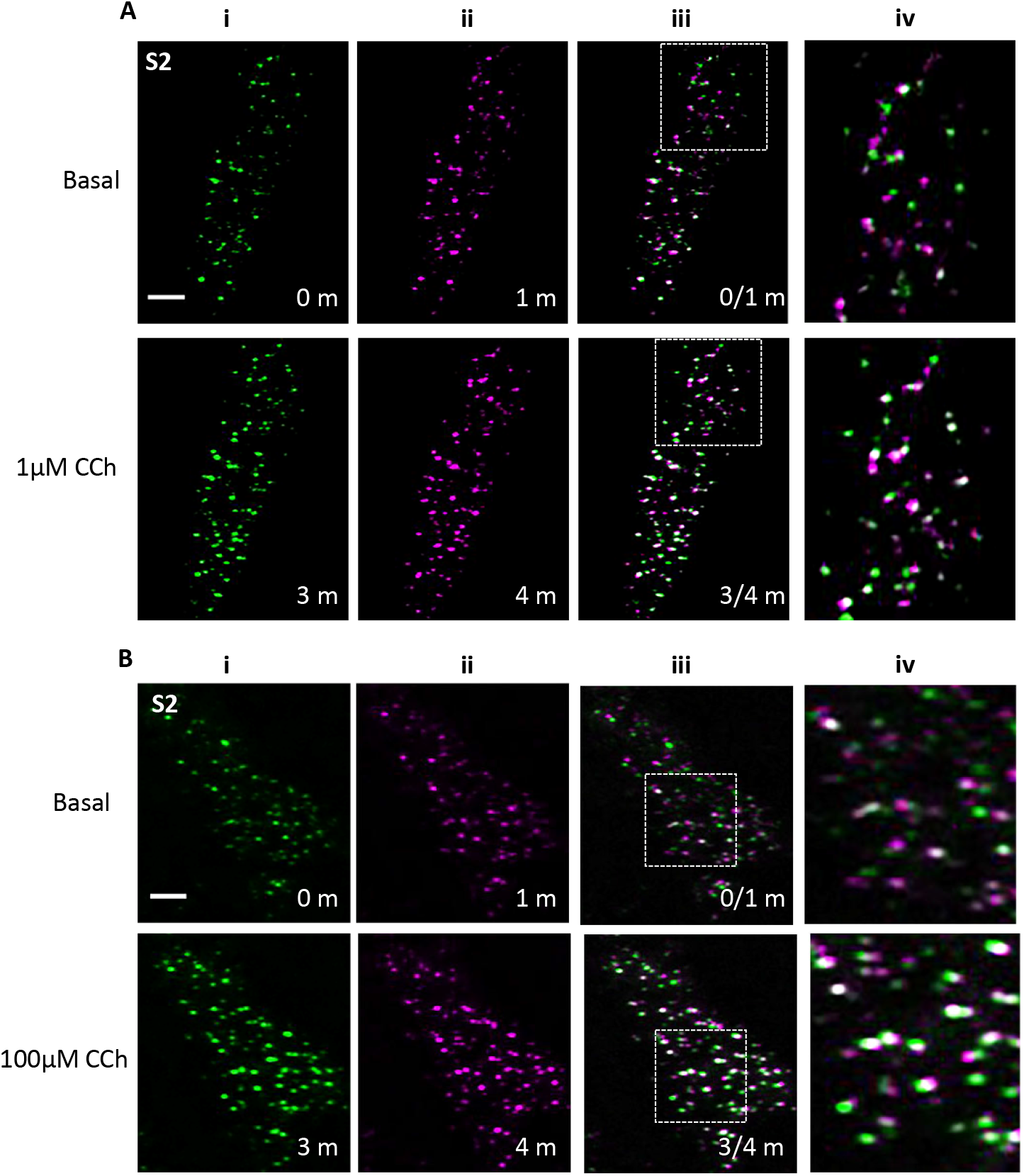
Mobility analysis of mVenus-STIM2 in STIM2-KI cells. (*A*) mVenus-STIM2 (mV- STIM2, labeled as S2) clusters in STIM2-KI cells without stimulation (basal condition, upper panels) and stimulated with 1µM CCh (lower panels). Images in each panel for (i, ii) were acquired at the start of experiment (0 m) and at the 1, 3 and 4 min time points (1, 3, and 4m). Images in (iii) were overlays of 0 and 1 min (0/1m), as well as 3 and 4 min (3/4m), time points. Images in (iv) are enlargements of the region marked by square in (iii). Overlapping clusters at the two time points appear white and indicate immobile mV- STIM2 clusters in basal condition and after CCh stimulation. (*B*) Images shown are in a similar order as in (*A*) except that cells were stimulated with 100µM CCh. All TIRFM images are representative of data obtained from 3 experiments. CCh was added at the 2 min time point for all experiments. Scale bar, 5µm.

**Fig. S3.**
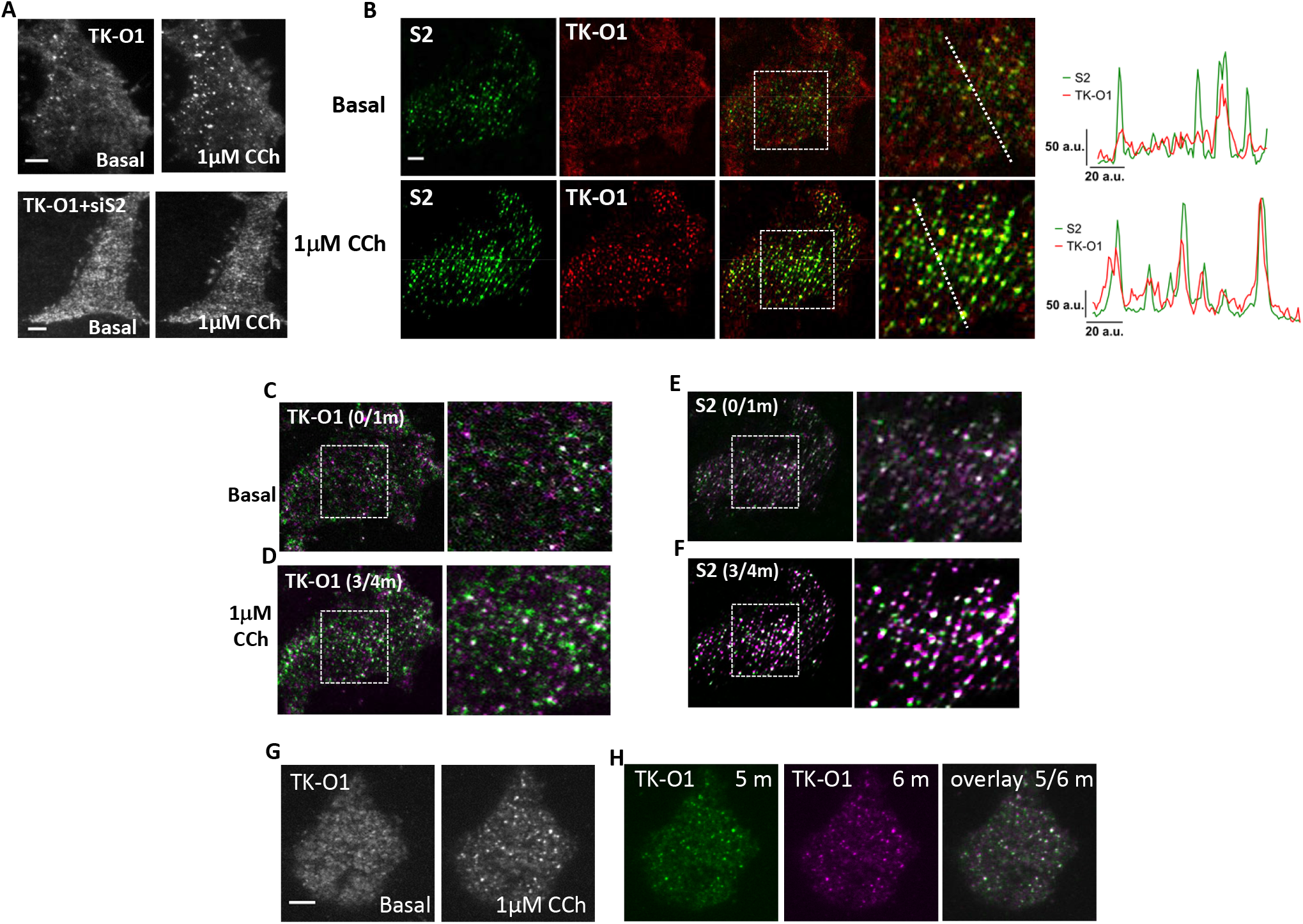
Analysis of TK-Orai1-CFP expression in HEK293 cells. (*A*) HEK293 cells showing TK-Orai1-CFP (TK-O1) without (upper panel) and with (siS2) siSTIM2 treatment, before (basal) and after 1μM CCh stimulation. (*B*) STIM2-KI cells expressing TK-Orai1-CFP (TK-O1) before (Basal; top) and after stimulation (1μM CCh; bottom). From left to right: mVenus-STIM2 (S2; Green); TK-O1 (Red); overlay of both proteins; enlarged area of overlay; and corresponding line scans. (*C-F*) of mV-STIM2 and TK-Orai1-CFP before (basal; 0/1m overlay) and after (3/4m overlay) 1μM CCh stimulation. Immobile clusters found at the same position in both frames appear as white. (*G*) HEK293 cells expressing TK-O1 during unstimulated (basal) conditions and after 1µM CCh stimulation, which is representative protein localization in majority of the cells (no preclusters). (*H*) TK-O1 clusters in wild-type HEK293 cells following stimulation with 1µM CCh. Images at the 5 and 6 min time points were pseudo-colored green (5m) and magenta (6m) were overlaid, with immobile clusters appearing white (5/6m). All TIRFM images are representative of data obtained from 3 experiments. CCh was added at the 2 min time point for all experiments. Scale bar, 5µm.

**Fig. S4.**
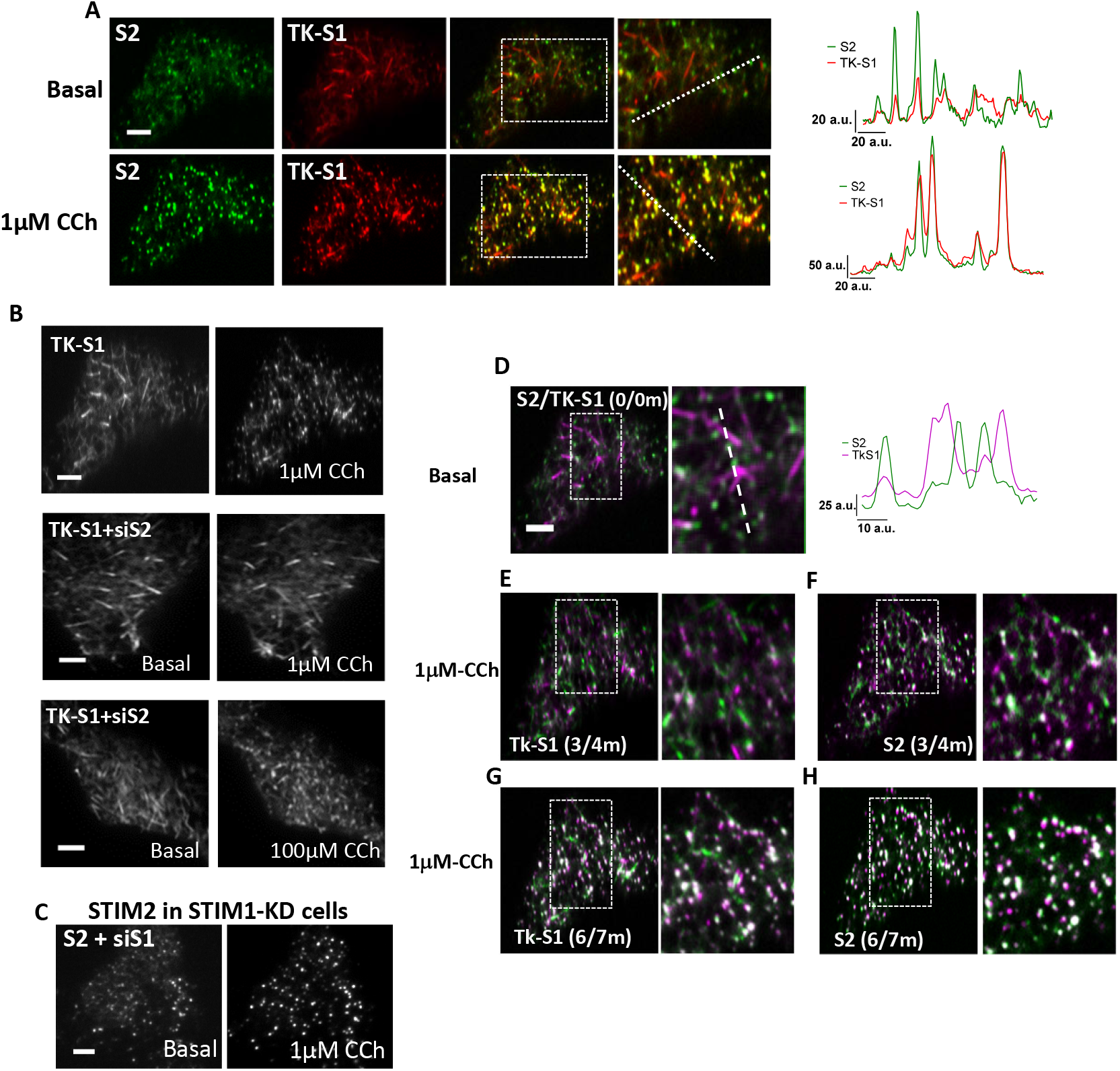
Analysis of TK-mCh-STIM1 expression in HEK293 cells. (*A*) STIM2-KI cell expressing TK-mCh-STIM1 (TK-S1), before (top) and after 1μM CCh stimulation (bottom). For each panel, the images are in the order of S2 (Green), TK-S1 (Red), overlay of both proteins, enlarged image and corresponding line-scans. (*B*) TK-mCh-STIM1 in HEK293 cells (upper panel) and siSTIM2-treated HEK293 cells (middle panel) before (basal) and after stimulation with 1μM CCh stimulation. Lower panel show representative cell treated with siSTIM2 in basal condition (lower left panel) and after 100 μM CCh (lower right panel). (*C*) Images show response to 1µM CCh in siSTIM1(siS1)-treated STIM2-KI cells. (*D*) Overlays of unstimulated STIM2-KI (S2, green) cells expressing TK-mCh-STIM1 (TK-S1, magenta) with the box showing the region used for scan analysis. (*E-H*) Overlay images of the same cell in (*B*) following addition of 1μM CCh: 2 min (4/5m overlay) and 4 min after stimulation (6/7m overlay) for (j, l) mV-STIM2 and (k, m) TK-STIM1-CFP. Immobile clusters found at the same position in both frames are white. All TIRFM images are representative of data obtained from 3 experiments. CCh was added at the 2 min time point for all experiments. Scale bar, 5µm.

**Fig. S5.**
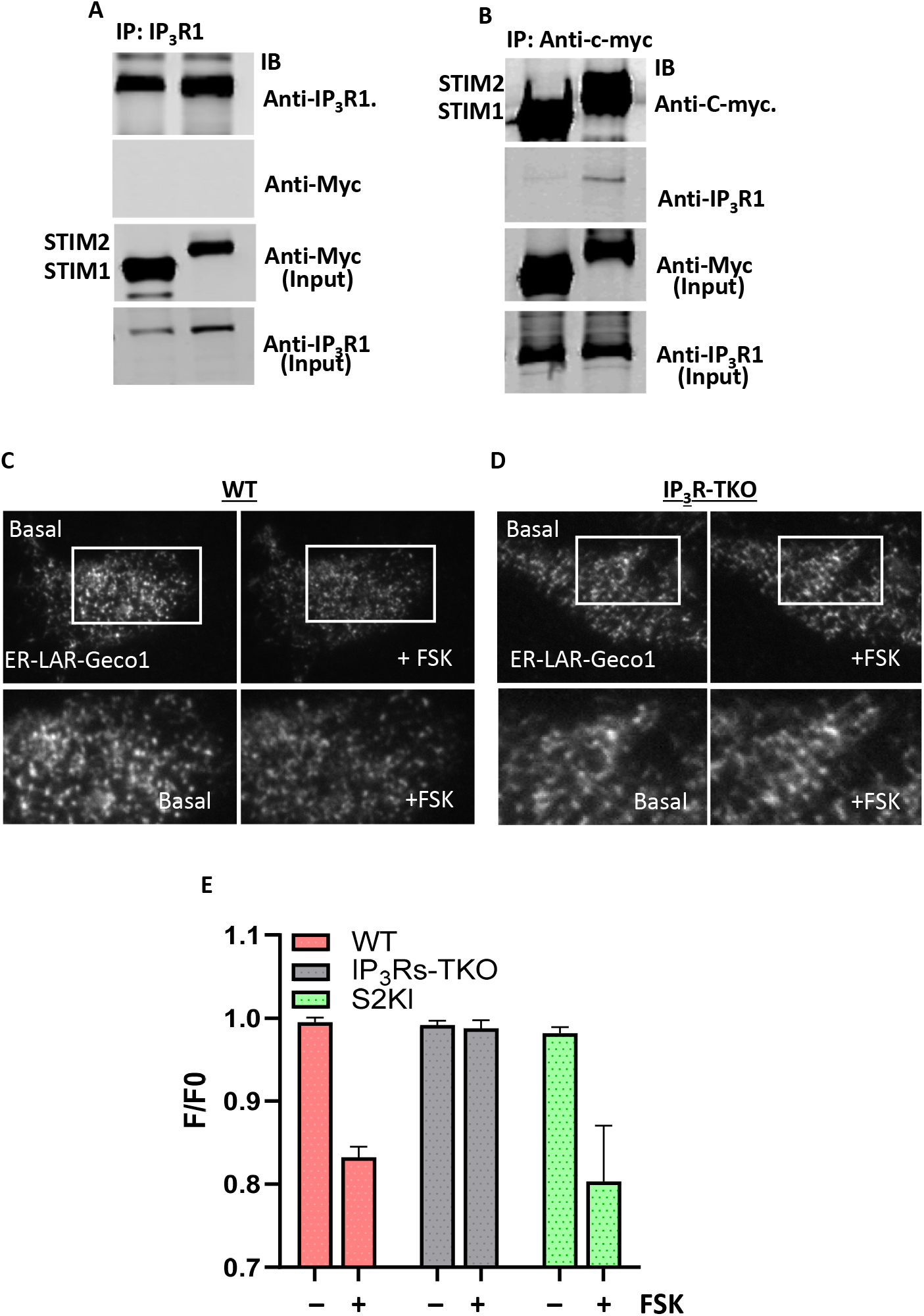
FSK-stimulated local [Ca^2+^]_ER_ depletion in IP_3_R-TKO cells and lack of Co-IP of STIM2 with IP_3_R1. (*A*) Immunoprecipitation (IP) of IP_3_R1 from HEK293 cells expressing Myc- STIM1 and Myc-STIM2. Immunoblotting (IB) was done using the anti-IP_3_R1 and anti-Myc antibody, respectively, for IP fractions (top and second blots) and input lysates (third and bottom blots). IB antibodies are shown next to each blot. (*B*) Reverse IP using the same cell lysate as in (*A*), but using anti-Myc antibody to pull down STIM1 and STIM2. The blots are representative of data obtained from 2 experiments. (*C*) HEK293 and IP_3_R-TKO cells expressing genetically encoded Ca^2+^ indicators, ER- Lar-GECO1 in basal (unstimulated condition), FSK (cells stimulated with Forskolin, 5µM). Enlargements of the region marked by square are shown for visible comparison. All images are TIRF micrographs and representative of data obtained from >3 experiments. FSK was added at the 2 min time point for all experiments. Scale bar, 5µm. (*D*) Bar graphs showing ER- Lar-GECO1 fluorescence (F/F_0_) before and after stimulation with 5µM FSK in wild-type (WT) HEK293, IP_3_R-TKO, and STIM2-KI (S2KI) cells. The graph shows averaged data (mean ± SEM) obtained from *n* = 16 Cells (WT), 13 Cells (IP_3_R-TKO), and 6 Cells (S2KI).

### Supplemental Video Legends

**Supplemental Video 1**: mVenus-STIM2 images in STIM2-KI cells were acquired every 10 s using TIRFM, video runs 5 frames/s. Frames 1-12 represent basal condition. 1µM CCh was added at frame 12.

**Supplemental Video 2**: mVenus-STIM2 images in STIM2-KI cells were acquired every 15 s using TIRFM, video runs 4 frames/s. Frames 1-8 represent basal condition with 1µM CCh addition at frame 9.

**Supplemental Video 3**: mVenus-STIM2images were acquired at every 10 s in siE-Syts2/3 treated STIM2-KI cells using TIRFM. Video runs 6 frames/s. Frames 1-12 represent basal condition, frames 12-36 with 1µM CCh, and frames 37-61 with 100µM CCh.

**Supplemental Video 4**: mVenus-STIM2 images were acquired every 10 second in siIP_3_Rs (all three subtypes were knocked down) treated STIM2-KI cells using TIRFM. Video runs 5 frames/s. Frames 1-12 represent basal condition and frames 12-36 with 100µM CCh.

**Supplemental Video 5**: mVenus-STIM2images were acquired every 10 s in STIM2-KI cells using TIRFM. Video runs 5 frames/s. Frames 1-6 represent basal condition and frames 7-31 with 5µM Forskolin.

**Supplemental Video 6**: TK-YFP-STIM2 expressed in HEK293 cells was imaged using TIRFM. Images were acquired at every 10 s. Video runs 5 frames/s. Frames 1-12 represent basal condition and frames 7-31 with 5µM Forskolin.

**Supplemental Video 7**: TK-YFP-STIM2 expressed in IP_3_R-TKO cells was imaged using TIRFM. Images were acquired at every 10 s. Video runs 5 frames/s. Frames 1-12 represent basal condition and frames 7-31 with 5µM Forskolin.

**Supplemental Video 8**: TK-YFP-STIM2 expressed in IP_3_R-TKO+IP_3_R1 cells was imaged using TIRFM. Images were acquired at every 10 s. Video runs 5 frames/s. Frames 1-12 represent basal condition and frames 7-31 with 5µM Forskolin.

**Supplemental Video 9:** TkS2 and TkS1N-S2C chimera (side by side) in unstimulated cells. Imaging was performed using TIRFM and each frame was acquired at every 10 s upto 2 min **Supplemental Video 10:** TkS1 and TkS2N-S1C chimera (side by side) in unstimulated cells. Imaging was performed using TIRFM and each frame was acquired at every 10 s upto 2 min.

